# Conservation and diversification of genes regulating brassinosteroid biosynthesis and signaling

**DOI:** 10.1101/2024.03.26.586792

**Authors:** Brian Zebosi, Erik Vollbrecht, Norman B. Best

## Abstract

Brassinosteroids (BRs) are important regulators that control myriad aspects of plant growth and development including biotic and abiotic stress responses, such that modulating BR homeostasis and signaling presents enormous opportunities for plant breeding and crop improvement. Enzymes and proteins involved in the biosynthesis and signaling of BRs are well understood from molecular genetics and phenotypic analysis in *Arabidopsis thaliana*; however, knowledge of molecular function of these genes in other plant species, especially cereal crop plants, is highly limited. In this manuscript, we comprehensively review functional studies of BR genes in Arabidopsis, maize, rice, Setaria, Brachypodium, and soybean to identify conserved and diversified functions across plant species, and to highlight cases where additional research is in order. We performed phylogenetic analysis of gene families known to be involved in biosynthesis and signaling of BRs and re-analyzed publicly available transcriptomic data. Gene trees coupled to expression data provide a useful guide to supplement future research on BRs in these important crop species, such as to allow researchers to identify genes to target through gene editing techniques to perform BR-related functional studies.

## Introduction

Brassinosteroids (BRs) are a class of plant-specific steroidal hormones. Their function in regulating plant growth and development has been studied in many different plant species. BRs, originally described as ‘brassins’, were first extracted from *Brassica napus* (rapeseed) pollen as a crude lipid extract and when mixed with lanolin resulted in strong elongation of internodes of a bean plant grown in the light (Mitchell et al., 1970). The structure of BRs, specifically brassinolide (BL), was determined in 1979 from an extraction of 500 lbs. of bee-collected rapeseed pollen (Grove et al., 1979). This was the first description of the growth-promoting BRs as being a class of polyhydroxyl lactones that commonly have a 5 α-cholestane skeleton, like that of animal steroids. BRs have been identified in green algae as well as bryophytes, moss, lycophytes, gymnosperms, and angiosperms (Yokota et al., 1987). They are present in all plant tissues and organs, but their highest abundance has been found in pollen and seeds (Bajguz and Tretyn, 2003).

BR function has been studied heavily in *Arabidopsis thaliana* (Arabidopsis) due to the development of many mutants in different biosynthetic enzymes and signaling and response components of BRs. Some work on BRs has been published in other model species; including *Zea mays* (maize), *Oryza sativa* (rice), and *Setaria viridis* (Setaria), *Brachypodium distachyon* (Brachypodium), and *Glycine max* (soybean); however, there are still many unknown gene functions across species due to an absence of mutant lines available. Investigating the conservation and/or diversification of these different gene families in eudicots and monocots is necessary to broadly understand BR gene functions across species. To that end, identifying gene homologs facilitates selecting reverse genetic targets to investigate these biological questions. To accomplish this, we conducted comprehensive phylogenetic analysis of encoded BR gene products across Arabidopsis, maize, rice, Brachypodium, soybean, and Setaria, re-analyzed publicly available transcriptomic experiments across multiple tissues in those species and presented those data together for BR biosynthesis, signaling, homeostasis, and response components. We selected genes based upon research findings that show the genes were involved in brassinosteroid biosynthesis, signaling, or catabolism in at least one of the presented species. To assist the reader, we have indicated in the text and figures an abbreviation of the specific genus and species we are describing before the gene name (i.e. *At* (Arabidopsis), *Zm* (maize), *Os* (rice), *Sv* (Setaria), *Bd* (Brachypodium), and *Gm* (soybean)).

### Brassinosteroid Independent Biosynthesis Pathway

Complex networks of biosynthetic pathways have been uncovered in plants for producing BRs. We focus in this article on the primary pathways leading to the sterols tracing through cycloartenol to active BRs. We describe this as the BR independent pathway, in that these mutant phenotypes cannot be recovered by exogenous BR treatment. Other reviews treat more comprehensively a broader range of possible pathways leading to BRs, e.g. including the mevalonate and non-mevalonate pathway branches upstream of sterols and the alternative pathway branch through cycloartanol and downstream of sterols (Bajguz et al., 2020).There are over 250 different sterol compounds in plants and the first dedicated compound in the biosynthesis of these phytosterols is squalene (Schaller, 2003; Schaller, 2004; Vriet et al., 2013).

Cycloartenol synthesis from squalene is a two-step reaction; the monooxygenation of squalene to squalene oxide/squalene epoxide by SQUALENE MONOOXYGENASES also known as SQUALENE EPOXIDASES (SQEs) and the cyclization of squalene oxide to either cycloartenol by CYCLOARTENOL SYNTHASE1 (CAS1) in plants and algae (Figure 1) or lanosterol by LANOSTEROL SYNTHASE (LSS) in animals and fungi (Rasbery et al., 2007; Desmond and Gribaldo, 2009; Laranjeira et al., 2015). SQEs are endoplasmic reticulum (ER)-localized and rate-limiting enzymes (Rasbery et al., 2007; Laranjeira et al., 2015) that are conserved across several eukaryotes with origins tracing back to Chlorophyta and charophytes (Desmond and Gribaldo, 2009) (Pollier et al., 2019). Mammals and fungi have a single SQE gene, whereas most higher plants contain multiple SQEs (Laranjeira et al., 2015; Pollier et al., 2019).

**Figure 1.**
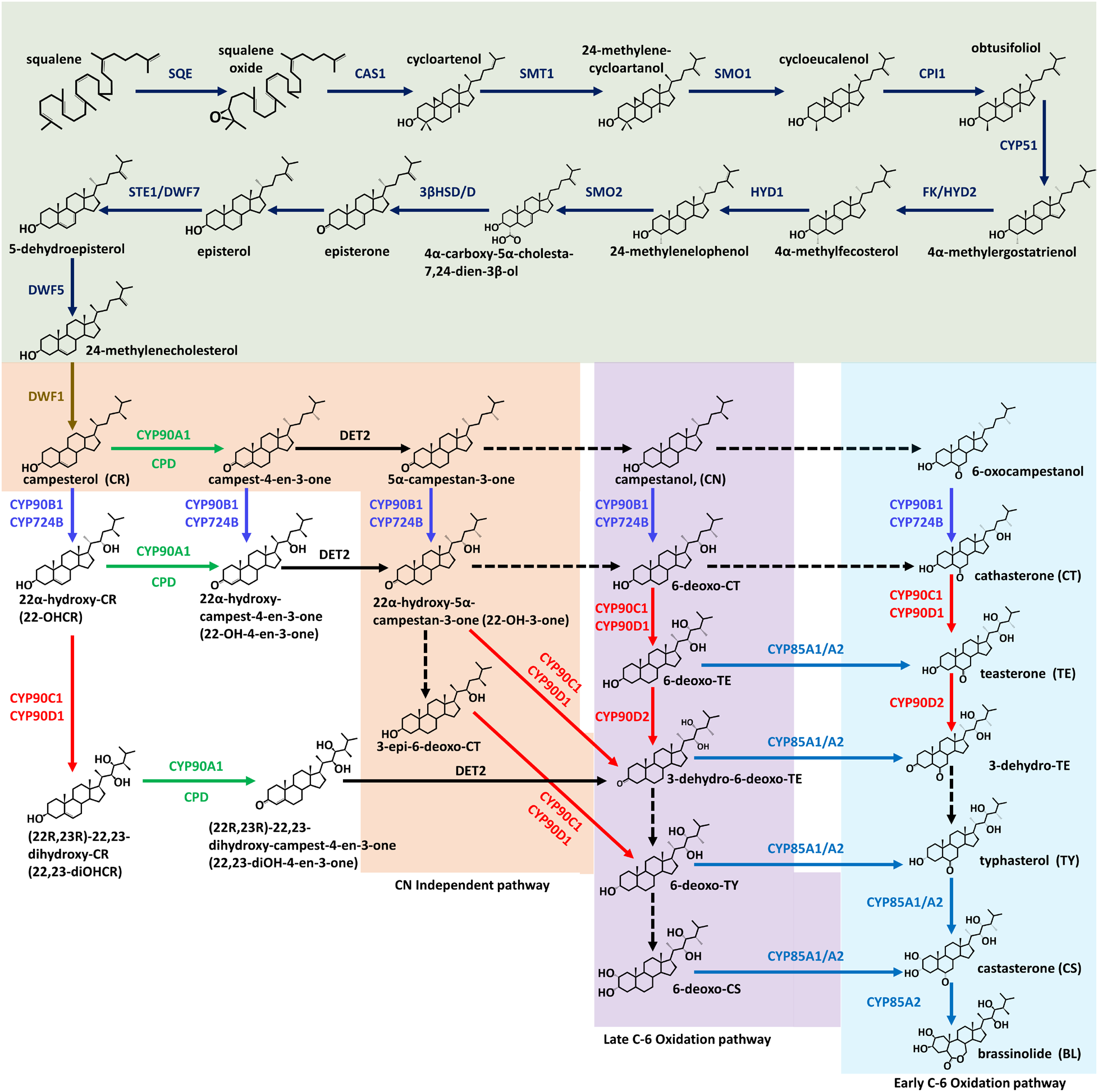
Phytosterol and brassinosteroid biosynthetic pathway. The phytosterol pathway is highlighted in green. The brassinosteroid early C-22 oxidation pathway is highlighted in orange, the late C-6 oxidation is highlighted in purple, and the early C-6 oxidation pathway is highlighted in blue. The chemical structure of the sterol intermediates is shown with the intermediate name written in black. Enzymes involved in the biosynthetic pathways are indicated above, below, and/or next to the arrows. Dashed arrow indicates multiple steps are possible. This pathway was modified from Ohnishi et al., 2012 and Kegg steroid biosynthesis. The pathway chemical structures were obtained from PubChem (Kim et al., 2023) and modified using the ChemDraw software.

Previous phylogenetic analyses (Rasbery et al., 2007) and our SQE gene tree (Figure 2) show that *SQEs* are single- or two-copy genes in most species, although six *SQE* family members exist in Arabidopsis (Figure 2A). The six Arabidopsis *SQE* genes clustered together in our analysis but were poorly resolved with respect to the *SQE*s of other species. *SQE2* and *SQE3* in Arabidopsis arose from a later duplication event (Posé et al., 2009; Laranjeira et al., 2015). Previous studies, in Arabidopsis have distinguished the *SQE-like* group (*AtSQE4*, *AtSQE5*, and *AtSQE6*) that resulted from a distinct duplication event (Figure 2A) (Posé et al., 2009; Laranjeira et al., 2015), from the other, “true” *AtSQE* genes. Interestingly, the “true” *AtSQEs* are phylogenetically closer to *SQE*s in humans and yeast than the *AtSQE*-like genes (Rasbery et al., 2007). In addition to sequence disparities, unlike SQE-like, the SQE proteins complemented the mutant defects of *ergosterol1* (*erg1*) in yeast (Rasbery et al., 2007; Posé et al., 2009; Laranjeira et al., 2015).

**Figure 2.**
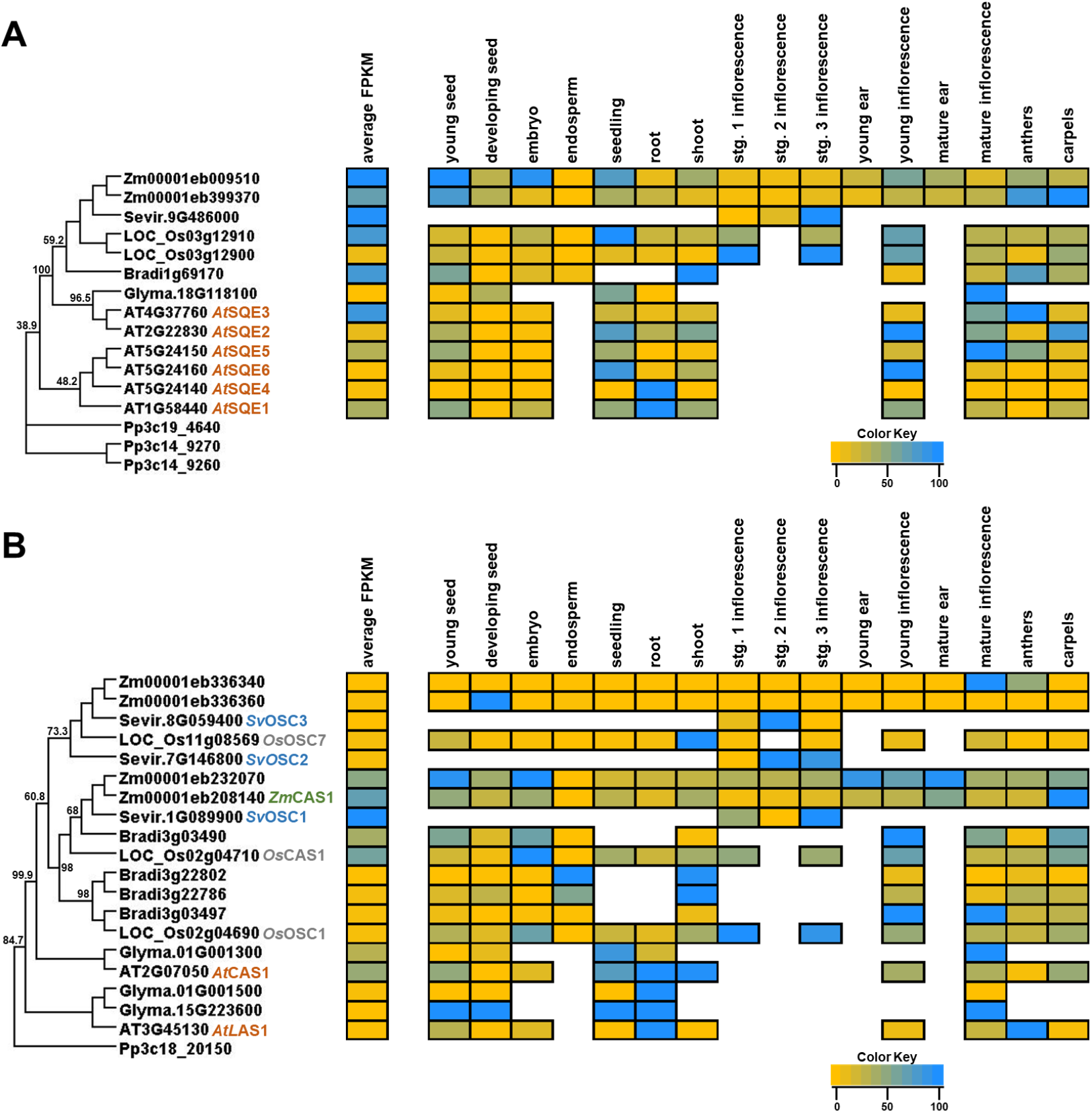
Phylogeny and transcript abundance across different tissues of SQUALENE EPOXIDASE (SQE) and CYCLOARTENOL SYNTHASE/LANOSTEROL SYNTHASE (CAS/LAS) phytosterol biosynthetic enzyme families. Maximum-approximate-likelihood phylogenetic trees of (A) SQE and (B) CAS/LAS amino acid sequences from maize (Zm), Setaria (Sevir), rice (Os), Arabidopsis (AT), soybean (Glyma), Brachypodium (Bradi), and Physcomitrella (Pp) represented as the outgroup. The relative TPM values from re-analyzed publicly available datasets from different developmental tissues are represented as a heatmap depicted next to each respective gene within the family. The yellow color indicates the lowest individual transcript abundance in each tissue and the dodgerblue color indicates the highest abundance in a given tissue. To compare transcript abundance across genes and species within the family, the average FPKM was determined across all tissues analyzed and is presented in the average FPKM column (left) with the same color distribution as previously described. See Supplemental Tables 1-4 for description of tissue types.

Mining of public expression data and published literature indicated overlapping and organ-specific expression patterns for *SQE* genes. In Arabidopsis, *SQE-like* genes were expressed at relatively low levels compared to “true” *SQE*s (Figure 2A) consistent with previous single-gene studies (Rasbery et al., 2007). *AtSQE3* is the most highly expressed overall among the *SQE*s, and *AtSQE1* and *AtSQE3* are expressed similarly in multiple tissues (Figure 2A; (Laranjeira et al., 2015)). In maize, two true *SQE*s have the highest average expression across tissues and are overlappingly expressed in young seeds; overall, differential expression of Zm00001eb399370 and Zm00001eb009510 suggests functional divergence after duplication. The Brachypodium and rice *SQE*s displayed overall similar expression patterns to the maize paralogs.

In contrast to the *AtSQE*-like gene family, the *AtSQEs* have been extensively studied and are implicated in plant growth and development. For instance, loss of function mutations in *AtSQE1* resulted in the accumulation of squalene and severe developmental defects such as reduced fertility and seed viability, aberrant root development with short and highly branched roots, and reduced plant height due to failed stem elongation (Rasbery et al., 2007; Posé et al., 2009; Laranjeira et al., 2015). While the *Atsqe2* and *Atsqe3* single and double mutants were not distinguishable from the wild type, the *Atsqe1; Atsqe3* double mutant displayed enhanced *Atsqe1* growth defects and embryo lethality, thus suggesting functional redundancy (Laranjeira et al., 2015). Importantly, these mutant developmental defects were not rescued by exogenous BR application implying that the SQEs contribute to other pathways apart from BR. Other homologs, including the Arabidopsis *SQE*-like genes and all *SQE* genes in soybean, Brachypodium, maize, rice, and Setaria, have not been functionally characterized.

#### Squalene oxide to cycloartenol or lanosterol

In the second reaction of sterol biosynthesis, squalene oxide is cyclized to cycloartenol and lanosterol by CYCLOARTENOL SYNTHASE1 (CAS1) and LANOSTEROL SYNTHASE (LAS), respectively (Babiychuk et al., 2008; Bach, 2016; Ohyama et al., 2009, Suzuki et al., 2006). The synthesis of lanosterol produces cholesterol, ergosterol, and phytosterol in mammals, fungi, and some plants, respectively (Kolesnikova et al., 2006; Suzuki et al., 2006). In higher plants, squalene oxide is predominantly cyclized to cycloartenol by CAS1, although eudicots also have LASs that metabolize squalene oxide to lanosterol (Kolesnikova et al., 2006; Suzuki et al., 2006) (Figure 2B). LASs and CASs share ∼65% amino acid identity supporting the proposition that LASs evolved from the ancient CASs after the monocot-dicot divergence.

Within the plant CAS1/LAS1 gene tree (Figure 2B) the CAS1-like clade is most predominant while consistent with other studies LAS1 genes are only found in eudicots, with two copies in soybean and one in Arabidopsis. CAS1 gene families have undergone differential expansion in the monocots examined. Across tissues, CAS1 clade members were expressed more highly than LAS1 genes on average (Figure 2B) and exhibited overlapping and tissue-specific expression patterns. For instance, monocot CAS1 genes, especially in maize and rice, were predominantly expressed in the shoot and inflorescences, whereas the dicot ones showed their highest expression in the roots. Within monocots, CAS1s displayed differential expression across tissues, consistent with sub- or neo-functionalization. In maize, Zm00001eb232070 showed higher expression than Zm00001eb208140 in most tissues except carpels (Figure 2B). Similarly, as reported in rice (Xue et al., 2012), in multiple tissues *OsCAS1* was more highly expressed than *OsOSC1*. Dicot LAS1 genes were expressed primarily in roots (Figure 2B; (Ohyama et al., 2009)).

Functional studies in Arabidopsis have highlighted the essential roles of CAS1 in plant growth and development. For example, *Atcas1* loss of function mutants were male gametophyte-lethal and exhibited pleiotropic developmental defects such as reduced plant growth, fused-multiple cotyledons, and arrested shoot apical meristem activity, albino shoots and inflorescences (Babiychuk et al., 2008; Go et al., 2012). Developmental roles of CAS1 in other species (soybeans, maize, rice, Seteria, and Brachypodium) remain unknown due to the lack of mutants. Gene editing approaches will likely be pivotal for generating the multiple mutant combinations needed to identify roles of CAS1 paralogs in these species.

Neither loss nor gain of *LAS1* function resulted in developmental defects or alteration in sterol profiles in Arabidopsis, suggesting cycloartenol as the primary and predominant pathway over lanosterol (Suzuki et al., 2006). Overexpression of LAS1 failed to rescue the *Atcas1* mutant phenotypes, confirming functional divergence between CAS1 and LAS1 (Suzuki et al., 2006; Ohyama et al., 2009). To date, no clear function of LAS1 has been reported; however, gene expression increases with treatment of methyl Jasmonate and *Pseudomonas* would suggest its involvement in plant defense responses (Kolesnikova et al., 2006; Ohyama et al., 2009). Genome editing would enable construction of double mutants to better understand the functional relationship between LAS1 and CAS1.

#### Cycloartenol to 24-methylenecycloartanol

In plants and fungi, phytosterols have extra alkyl groups at C-24; this alkylation is catalyzed by STEROL C-24 METHYLTRANSFERASEs (SMTs) via S-adenosylmethionine (SAM)-dependent transmethylation (Diener et al., 2000; Holmberg et al., 2002). Contrary to fungi, with a single methyl group at C-24, higher plants undergo sequential C-24 methylation reactions catalyzed by distinct *SMT* genes. The first C-24 methylation is executed by SMT1s that are conserved across several kingdoms and convert cycloartenol to 24-methylene-cycloartenol, that serves as precursor of campesterol (CR, a 24-methyl sterol) and sitosterol (a 24-ethyl structural sterol). Sequentially, SMT2 and/or SMT3 enzymes that diverged from the ancestral *SMT*s in Charophytes catalyze the second methylation at the side chain of 24-methylenelophenol to produce 24-ethylidenelophenol, a vital intermediate for sitosterol synthesis (Carland et al., 2010; Chen et al., 2018; Gallo et al., 2020) (Haubrich et al., 2015; Nakamoto et al., 2015). In addition to this functional specialization, SMT1 and SMT2/SMT3 have contrasting roles in balancing cholesterol with other sterols and regulating the ratio of campesterol to sitosterol (Diener et al., 2000; Carland et al., 2002).

Phylogenetic analysis showed expansion of both the *SMT1* and the *SMT2*/*SMT3* gene families (Figure 3A). Across the clades, *SMT1* genes showed higher expression than the *SMT2/SMT3*s in most tissues. However, within clades, expression is diverse across tissues. In Arabidopsis, *AtSMT1* is primarily expressed in developing embryos, shoot, and root. Whereas, *AtSMT2* and *AtSMT3* are moderately expressed in most tissues except in young seeds and roots (Figure 3A; (Diener et al., 2000)). Similar to Arabidopsis, *SMT1* genes in Brachypodium, rice and maize are extensively expressed in most tissues, while the *SMT2/SMT3* members in these species, are primarily expressed in the inflorescences. The differential gene expression suggests functional diversification between *SMT1* and *SMT2/SMT3*.

**Figure 3.**
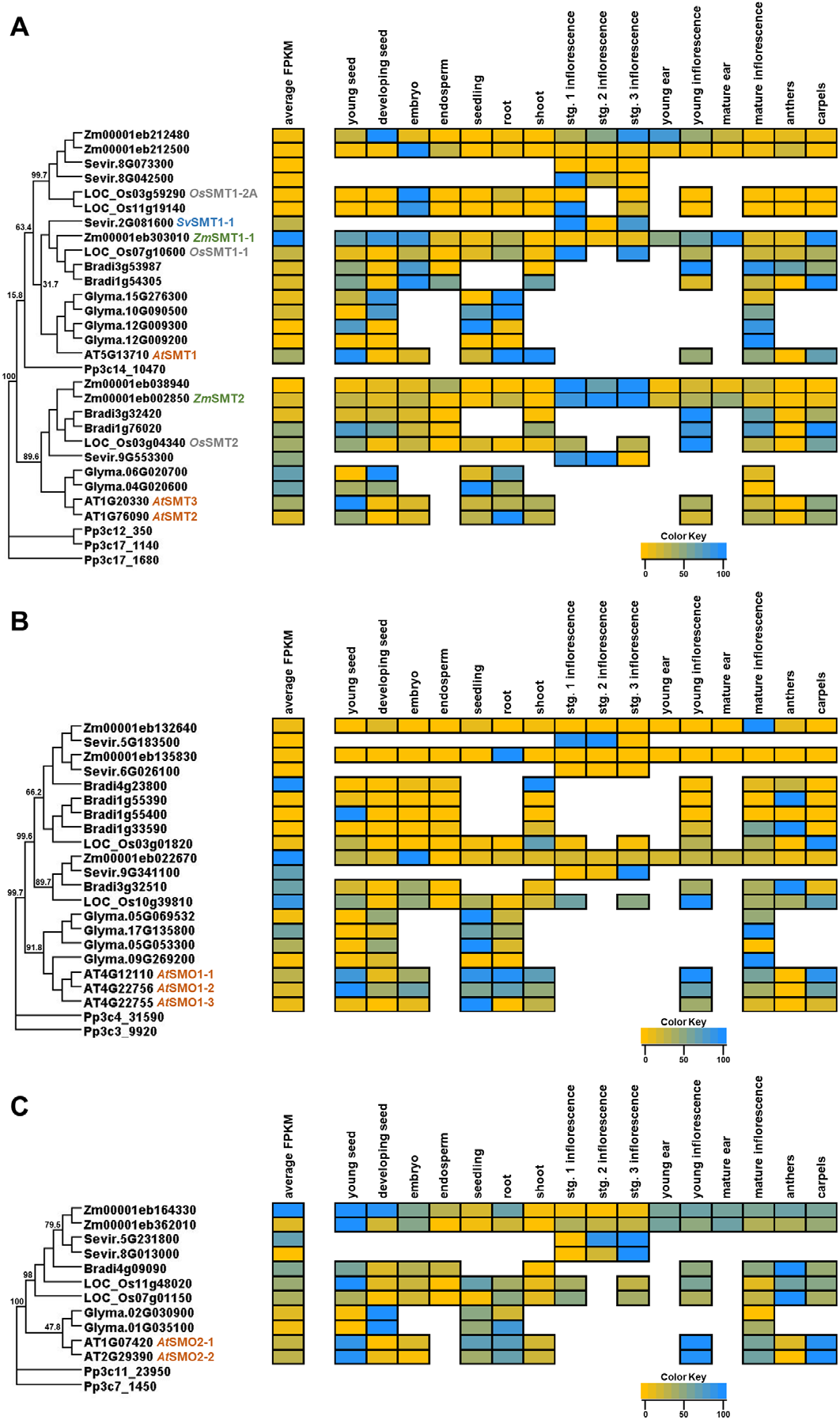
Phylogeny and transcript abundance across different tissues of STEROL METHYLTRANSFERASE (SMT), STEROL 4α-METHYL OXIDASE1 (SMO1), and STEROL 4α-METHYL OXIDASE (SMO2) phytosterol biosynthetic enzyme families. Maximum-approximate-likelihood phylogenetic trees of (A) SMT, (B) SMO1, and (C) SMO2 amino acid sequences from maize (Zm), Setaria (Sevir), rice (Os), Arabidopsis (AT), soybean (Glyma), Brachypodium (Bradi), and Physcomitrella (Pp) represented as the outgroup. The relative TPM values from re-analyzed publicly available datasets from different developmental tissues are represented as a heatmap depicted next to each respective gene within the family. The yellow color indicates the lowest individual transcript abundance in each tissue and the dodgerblue color indicates the highest abundance in a given tissue. To compare transcript abundance across genes and species within the family, the average FPKM was determined across all tissues analyzed and is presented in the average FPKM column (left) with the same color distribution as previously described. See Supplemental Tables 1-4 for description of tissue types.

The *SMT1* and *SMT2/SMT3* gene families in Arabidopsis have been shown to regulate several developmental processes such as embryo morphogenesis, cotyledon vein patterning, apical dominance, auxin response, and reproductive and root development (Carland et al., 2010; Nakamoto et al., 2015). The *Atsmt1* mutants exhibit several pleiotropic defects, including reduced plant stature with low fertility and aberrant embryo morphogenesis with aborted and dulled embryos (Diener et al., 2000). Consistent with expression data, *AtSMT2* and *AtSMT3* function collectively to regulate plant development. While single mutants were affected weakly or not at all, the *Atsmt2*;*Atsmt3* double mutant displayed synergistically enhanced developmental defects such as extreme dwarfism, self-sterility, irregular vasculature, aberrant root development, deformed cell shape, and disorganized tubulin orientation (Carland et al., 2010). In addition to reduced sitosterol and CR, these mutants also had impaired gravitropism with aberrant localization of auxin efflux carriers (Carland et al., 2010; Nakamoto et al., 2015).

By contrast to the extensive functional studies in Arabidopsis, no similar studies of *SMT1* or *SMT2*/*SMT3* have been reported in Brachypodium, Seteria, soybean, rice, or maize. Functional studies will be necessary to determine SMT roles in these species.

#### 24-methylene-cycloartanol to cycloeucalenol and 24-methylenelophenol to 4α-carboxy-5α-cholesta-7,24-dien-3β-ol

Conversion of (firstly) 24-methylenecycloartenol into cycloeucalenol and (secondly) of 24-methylenelophenol into episterol (Figure 1) is catalyzed by two independent STEROL C-4 METHYL OXIDASES (SMOs) catalyzing non-sequential C-4 demethylation reactions (Figure 1) unlike in fungi and animals, where a single SMO executes the two C-4 demethylation reactions sequentially (Lung et al., 2017; Song et al., 2019) (Sonawane et al., 2016; Song et al., 2019). The first and second demethylation reactions are catalyzed by SMO1 and SMO2, respectively (Sonawane et al., 2016; Song et al., 2019).

The *SMO1* gene family (Figure 3B) has between two and five genes per species examined. In Arabidopsis, *AtSMO1-1* and *AtSMO1-2* were widely and jointly expressed in most tissues, especially young seeds, inflorescences, and roots (Figure 3B; (Song et al., 2019)). *SMO1* genes in maize and Brachypodium showed expression patterns suggesting specialization across tissues such as embryos, young inflorescences, and mature inflorescences. In rice, two genes, LOC_Os10g39810 and LOC_Os03g01820, showed differential expression with LOC_Os03g01820 more widely expressed. In soybean, SMO1s were ubiquitously and moderately expressed across tissues. In the *SMO2* family (Figure 3C), Brachypodium has a single-copy gene, and the other species have two *SMO2*s. *SMO2* genes showed overall broader tissue expression patterns than their *SMO1* paralogs, and the degree of overlapping expression within species suggests a corresponding functional redundancy.

Functional characterization is restricted to Arabidopsis, where *SMO1* and *SMO2* single mutants had no obvious developmental defects, whereas both *Atsmo1-1*;*Atsmo1-2* and *Atsmo2-1;Atsmo2-2* double-mutants were embryo lethal with aborted shoot apical meristems with aberrant cell division and suppressed cytokinin levels (Song et al., 2019; Zhang et al., 2016). These double mutant combinations resembled auxin-defective mutants with accumulated 4α- methyl-sterols and auxin levels and impaired auxin transport (Zhang et al., 2016; Song et al., 2019). The double mutant embryo lethality was partially rescued by auxin or cytokinin treatment and *AtYUCCA9* overexpression (Zhang et al., 2016; Song et al., 2019). Thus, *SMO1*s and *SMO2*s are vital for embryogenesis and regulating auxin production and transport (Zhang et al., 2016; Song et al., 2019). Functional characterization in other species is required to examine these genes’ broader functional diversification or conservation.

#### Cycloeucalenol to 4α–methylergostatrienol

CYCLOPROPYLSTEROL ISOMERASE1 (CPI1) and OBTUSIFOLIOL 14α-DEMTHYLASE (CYP51G1) catalyze the two-step conversion of cycloeucalenol to 4α– methylergostatrienol (Figure 1) (Men et al., 2008). CPI1 catalyzes the isomerization of cycloeucalenol to obtusifoliol and it is restricted to land plants, as fungi have no CPI1 homolog (Lovato et al., 2000; Sonawane et al., 2016). CPI1 is encoded by a single gene in Arabidopsis, Brachypodium, maize, Setaria, and rice, and two duplicate genes in soybean (Figure 4A). *CPI1* genes showed distinct tissue-specific expression patterns across species, suggesting corresponding species-specific sterol utilization. Functional studies indicate that like the SMT1, SMT2/SMT3, SMO1 and SMO2 enzymes, CPI1 regulates several plant growth and development aspects such as embryo morphogenesis, and root development and auxin distribution (Men et al., 2008; Wang et al., 2021). The *Atcpi1* mutants were semi-sterile dwarf plants with short roots and impaired cell expansion and aberrant root gravitropism (Men et al., 2008), and impeded auxin transporter localization (Men et al., 2008; Wang et al., 2021). There were no reported functional studies of *CPI1* genes in monocots and soybean.

**Figure 4.**
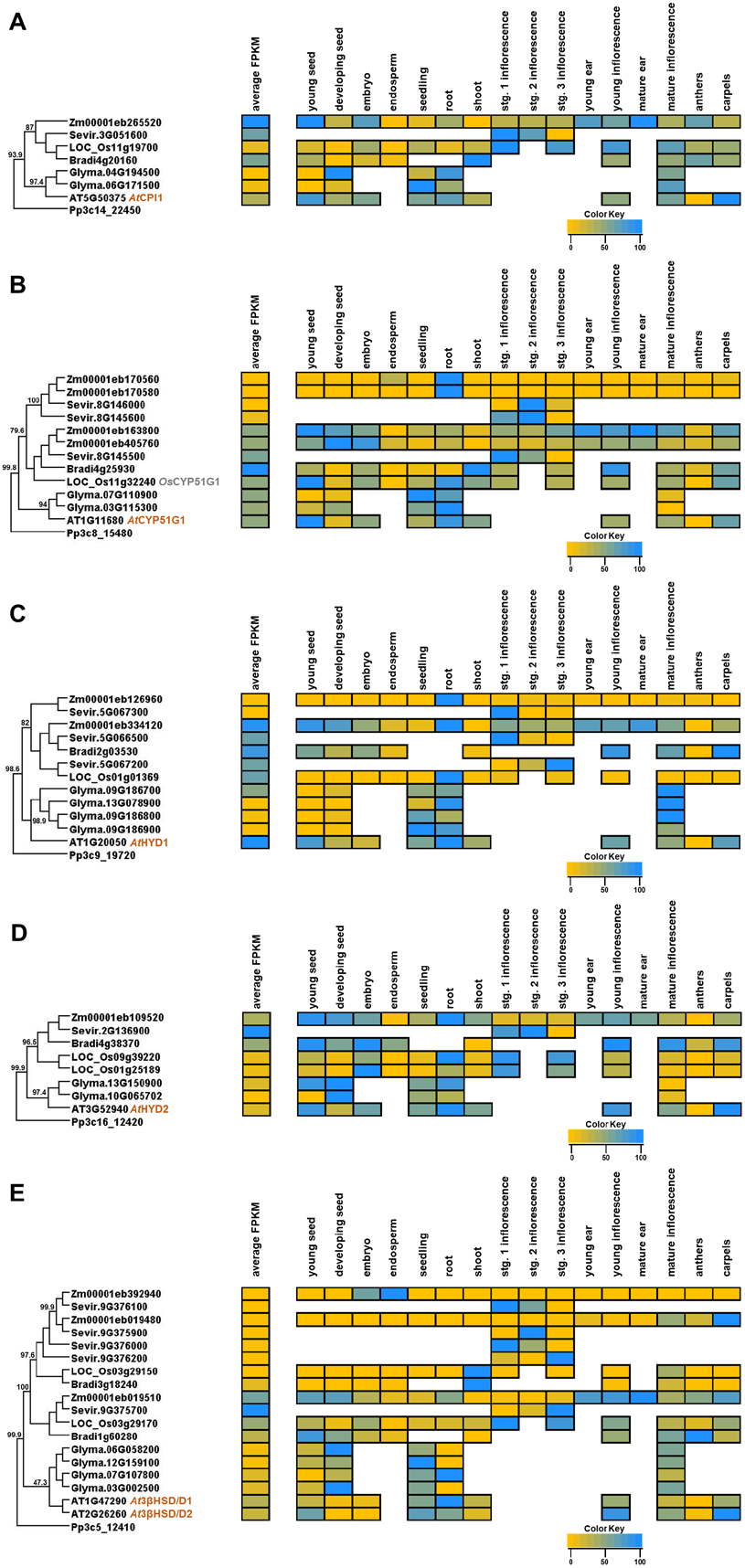
Phylogeny and transcript abundance across different tissues of CYCLOPROPYL ISOMERASE (CPI), CYTOCHROME P450 51G1 (CYP51G1), HYDRA1 (HYD1), HYDRA2 (HYD2), and 3β-HYDROXYSTEROID DEHYDROGENASE/C4 DECARBOXYLASE (3βHSD/D) phytosterol biosynthetic enzyme families. Maximum-approximate-likelihood phylogenetic trees of (A) CPI, (B) CYP51G1, (C) HYD1 and HYD2, and (D) 3βHSD/D amino acid sequences from maize (Zm), Setaria (Sevir), rice (Os), Arabidopsis (AT), soybean (Glyma), Brachypodium (Bradi), and Physcomitrella (Pp) represented as the outgroup. The relative TPM values from re-analyzed publicly available datasets from different developmental tissues are represented as a heatmap depicted next to each respective gene within the family. The yellow color indicates the lowest individual transcript abundance in each tissue and the dodgerblue color indicates the highest abundance in a given tissue. To compare transcript abundance across genes and species within the family, the average FPKM was determined across all tissues analyzed and is presented in the average FPKM column (left) with the same color distribution as previously described. See Supplemental Tables 1-4 for description of tissue types.

The second enzyme, CYP51G1, a cytochrome P450 (CYP) monooxygenase, is encoded by *CYP51G* gene family members and catalyzes the production of 4α–methylergostatrienol from obtusifoliol through C-14 demethylation (Kushiro et al., 2001). CYP51G1 is highly conserved in the animal, fungi, and plant kingdoms, and CYP51s utilize different sterol substrates across different kingdoms; for instance, higher plants use obtusifoliol to produce the precursors of BRs and structural sterols whereas animals and fungi use lanosterol to synthesize cholesterol and ergosterol, respectively (Debeljak et al., 2003; Kim et al., 2005; O’Brien et al., 2005; Geisler et al., 2013). In contrast with a single CYP51 copy in yeast and mammals, some plants have several functional CYP51s due to multiple duplication events (O’Brien et al., 2005; Lepesheva and Waterman, 2007). Our phylogenetic analysis (Figure 4B) indicated four canonical *CYP51* genes in maize; three in Setaria; two in soybean; and one in rice, Brachypodium and Arabidopsis. Notably, previous studies have indicated that the Arabidopsis and rice genomes have other *CYP51* genes, but only one is a functional *CYP51G1* (Søren et al., 2011). In rice, the *CYP51* family has 12 members, three *OsCYP51G*s (*1*,*3*,*4*) and nine *OsCYP51H*s (1-9), but only *OsCYP51G1* directly encodes a functional obtusifoliol 14α-demethylase (Inagaki et al., 2011; Xia et al., 2015; Jiao et al., 2020).

Investigating the expression patterns of CYP51G1 transcripts among and within species indicated that in general these genes are highly expressed in several tissues (Figure 4B). In contrast, low expression of maize Zm00001eb170560 and Zm00001eb170580 could imply functional diversification or reflect pseudogene status. Soybean *CYP51G1*s are primarily expressed in seedlings and roots while Bradi4g25930 is primarily expressed in young seeds and young inflorescences. In rice, *OsCYP51G1* show highest expression in shoots and roots, whereas *AtCYP51G1* was in young seeds and roots.

Functional characterization of *CYP51G1* genes in Arabidopsis and rice indicated that CYP51G1s are involved in shoot and embryonic development, anther and pollen development, reproductive heading, seed production, phytosterol biosynthesis, and auxin-ethylene signaling regulation (Kim et al., 2005; Jiao et al., 2020). In Arabidopsis, the *Atcyp51g1* mutants showed stunted growth with seedling lethality and sterility (Kushiro et al., 2001; Kim et al., 2005). The *Atcyp51g1* mutants also had reduced cotyledons and hypocotyls with defective cell expansion (Kim et al., 2005). The *Atcyp51g1* mutants were insensitive to exogenous BR treatment and accumulated obtusifoliol but had decreased CR and sitosterol levels (Kim et al., 2005).

In rice, loss of function mutants of *OsCYP51G1* exhibited similar growth defects as *Atcyp51g1*. In addition, the *Oscyp51g1* mutants had delayed flowering with abnormal anther development and pollen sterility (Jiao et al., 2020). *Os*CYP51G1 overexpression lines (*Os*CYP51G1-OE) had increased grain weight and more seeds per panicle, highlighting the potential of *Os*CYP51G1 in crop yield improvement (Jiao et al., 2020). Initial studies emphasized that *Os*CYP51G3 and *Os*CYP51H3 are phylogenetically distant from *Os*CYP51G1 (Nelson et al., 2004; Inagaki et al., 2011), but it was subsequently shown mutants of *Oscyp51g3* and *Oscyp51h3* displayed growth defects similar to those of *Oscyp51g1* mutants (Xia et al., 2015; Jiao et al., 2020). Thus, it is hypothesized that *Os*CYP51G1, *Os*CYP51G3, and *Os*CYP51H3 function collectively to regulate phytosterol and BR biosynthesis and plant growth and development. Double and triple mutants should be produced to address this claim. Due to the lack of mutants in Brachypodium, maize, soybean, and Setaria, the function of these species’ CYP51G1s remain to be investigated.

#### 4α–methylergostatrienol to 24-methylenelophenol

The enzymatic conversion of 4α–methylergostatrienol to 24-methylenelophenol occurs in two significant sequential reactions; first, the C14 reduction of 4α–Methylergostatrienol to 4α- Methyl-fecosterol by sterol C-14 reductase, encoded by *HYDRA2* (*HYD2*) in Arabidopsis and *ERG24* in yeast (Jang et al., 2000; Schrick et al., 2000); and then second, isomerization of 4α- methyl-fecosterol to 24-methylenelophenol by C-8,7 sterol isomerase, encoded by *HYD1* and *ERG2* in Arabidopsis and yeast, respectively (Souter et al., 2002). Sequence similarity and complementation experiments indicated that C-14 sterol reductase and C-8,7 sterol isomerase are conserved across plants, fungi, and animals (Jang et al., 2000; Pierre, 2002; Souter et al., 2002; Pullen et al., 2010). Our evolutionary analysis resulted in two separate phylogenetic trees; *HYD1* and *HYD2* (Figure 4C and 4D). The *HYD1* gene is single-copy in Arabidopsis, Brachypodium, and rice but shows different duplication patterns among soybean, Setaria, and maize (Figure 4C). The *HYD2* genes on the other hand, were strictly single- or two-copy with only intra-specific duplications (Figure 4D). Despite these evolutionary disparities HYD1 and HYD2 family members exhibited some similar expression patterns, such as overlapping and highest expression in roots, suggesting the possibility of some functional conservation between gene families and across species.

Functional roles of *HYD1* and *HYD2* in plant growth and development have been well characterized through mutant analysis and expression studies only in Arabidopsis. Loss of function mutants indicated that these genes are jointly implicated in embryonic cell patterning, vascular tissue development, auxin, and ethylene signaling (Souter et al., 2002). The *Athyd1* and *Athyd2* single mutants have dwarf plant stature with abnormal cell division, defective apical-basal radial patterns, and disorganized vascular patterns (Jang et al., 2000; Souter et al., 2002; Short et al., 2018). In addition, *Athyd1* and *Athyd2* mutants displayed increased ethylene response, reduced expression of auxin regulated genes (*IAA1* and *IAA2*), and aberrant polar localization of auxin transporters (PIN1 and PIN2) (Short et al., 2018); exogenous application of BR did not rescue the mutant growth defects, thus suggesting that *At*HYDs may function in the production of other phytohormones such as auxin, ethylene, and BR (Pullen et al., 2010). In monocots and soybean, the *HYD* genes remain uncharacterized, which should be a focus for future research.

#### 4α-carboxy-5α-cholesta-7,24-dien-3β-ol to episterol

The synthesis of episterol from 4α-carboxy-5α-cholesta-7,24-dien-3β-ol occurs in two successive reactions; the oxidative decarboxylation of 4α-carboxy-5α-cholesta-7,24-dien-3β-ol to episterone then the reduction of episterone to episterol, which is catalyzed by 3β- HYDROXYSTEROID DEHYDROGENASE/C4 DEXCARBOXYLASE (3βHSD/D) (Rahier et al., 2006) and a 3-keto steroid reductase (Gachotte et al., 1999), respectively. The 3βHSD/Ds belong to the short-chain alcohol dehydrogenase/reductase (SDR) family and are conserved in eukaryotes and function in ergosterol, cholesterol, and phytosterol biosynthesis in humans, fungi, and plants, respectively (König et al., 2000; Simard et al., 2005; Helliwell et al., 2015). The 3βHSD/D protein is encoded by *ERG26* in yeast and *3βHSD/D* in Arabidopsis (Rahier et al., 2006). Conversion of episterone to episterol requires a 3-keto reductase, however this enzyme has only been identified in yeast and no close homologs in plants were identified (Gachotte et al., 1999; Desmond and Gribaldo, 2009). Further studies are necessary to identify the functional enzyme responsible for this reaction in plants.

To examine the evolutionary relationship of *3βHSD/D* in plants, we constructed a gene tree that revealed that the 3βHSD/D is encoded by several *3βHSD/D* copies depending on the species (Figure 4E). There are five copies in Setaria, four in soybean, three in maize, and two in each of Arabidopsis, Brachypodium, and rice. Mined expression data indicated that one gene member was predominantly expressed in maize (Zm00001eb019510), in Brachypodium (Bradi1g60280), and in rice (LOC_Os03g29170), with additional tissue-specificity in each species. Family member expression in Arabidopsis, soybean, and Setaria was more overlapping, suggesting greater functional redundancy may exist in these species.

Functional characterization studies in Arabidopsis have implicated 3βHSD/Ds in several aspects of plant growth and development including regulation of auxin movement (Rahier et al., 2006). Neither the single mutants (*At3βhsd/d1* or *At3βhsd/d2*) nor the double mutant (*At3βhsd/d1*; *At3βhsd/d2*) displayed obvious growth defects (Kim et al., 2012). However, a line overexpressing both *At*3βHSD/D1 and *At*3βHSD/D2 had compact plant stature, wrinkled leaves, and a small inflorescence with hampered auxin localization. These growth defects were unaltered with BL treatment (Kim et al., 2012), though rescue was not attempted with BR inhibitors. To date, no *3βHSD/D*s have been characterized in monocots or soybean.

For the sterol biosynthesis genes discussed thus far in producing episterone from squalene, none of the mutant phenotypes have been reported to be rescued by exogenous BR application. By contrast, mutants for the next set of enzymes have growth and development phenotypes that, as a rule, are rescued by exogenous BR application. Thus, these following steps are required exclusively to produce BR, and together, they have been referred to in the literature as the “brassinosteroid-dependent” segment of the BR biosynthesis pathway (Clouse, 2002).

### Brassinosteroid Dependent Biosynthesis Pathway

#### Episterol to campesterol

After sequential reactions, including a C-4 demethylation catalyzed by SMO2 (discussed above) to produce episterol from 24-methylenelophenol, the start of the BR-dependent pathway is marked by the episterol to CR steps that occur in three successive reactions; first, the C-5 desaturation of episterol to 5-dehydro-episterol occurs by the activity of a delta-7 sterol C-5 desaturase encoded by *DWARF7*/*STEROL1* (*DWF7*/*STE1*) (Cheon et al., 2010; Silvestro et al., 2013). Secondly, sterol delta-5-dehydro-episterol conversion to 24-methylene-cholesterol via reduction is catalyzed by a sterol delta-7 reductase and encoded by *DWARF5* (*DWF5*) (Choe et al., 2000; Silvestro et al., 2013; Inoue et al., 2017). Thirdly, the C-24 reduction of 24-methylene-cholesterol by sterol C-24 reductase encoded by *DWARF1* (*DWF1*) (Choe et al., 1999) yields CR, a precursor sterol that leads to the genesis of the BR biosynthesis pathway (Choe et al., 1999; Silvestro et al., 2013; Youn et al., 2018).

These enzymes, DWF7, DWF5, and DWF1, also function successively outside of BR biosynthesis in the sitosterol-stigmasterol biosynthetic pathway by catalyzing the desaturation of delta-7-avenasterol to 5-dehydro-avenasterol (Clouse, 2002), delta-7 reduction of 5-dehydro-avenasterol into isofucosterol (Choe et al., 2000) and reduction of isofucosterol to sitosterol (Clouse, 2002), respectively. These reactions are parallel to the conversion of episterol to CR (Clouse, 2002). Stigmasterol has been shown to function in cell membrane integrity and physiology and act as a potential signaling molecule involved in gravitropism and defense mechanisms (Griebel and Zeier, 2010; Aboobucker and Suza, 2019). Thus, mutations in *DWF1*, *DWF5*, or *DWF7* result in CR and sitosterol deficiency, associated with both altered plant development and defective membrane structure (Clouse, 2002). Evolutionarily, DWF1, DWF5, and DWF7 (Figure 5A-C) are highly conserved enzymes across the plant and fungal kingdoms (Sonawane et al., 2016), suggesting that their genes may have arisen early in eukaryotic evolution (Vriet et al., 2015). Interestingly, these genes split into separate clades with different outgroups (Figure 5A-C), indicating multiple duplication rounds resulting in enzymes with specific functions (Desmond and Gribaldo, 2009; Sonawane et al., 2016). Their gene family sizes vary, with only *DWF5* showing no evidence of gene duplication in the species reviewed here (Figure 5A-C).

**Figure 5.**
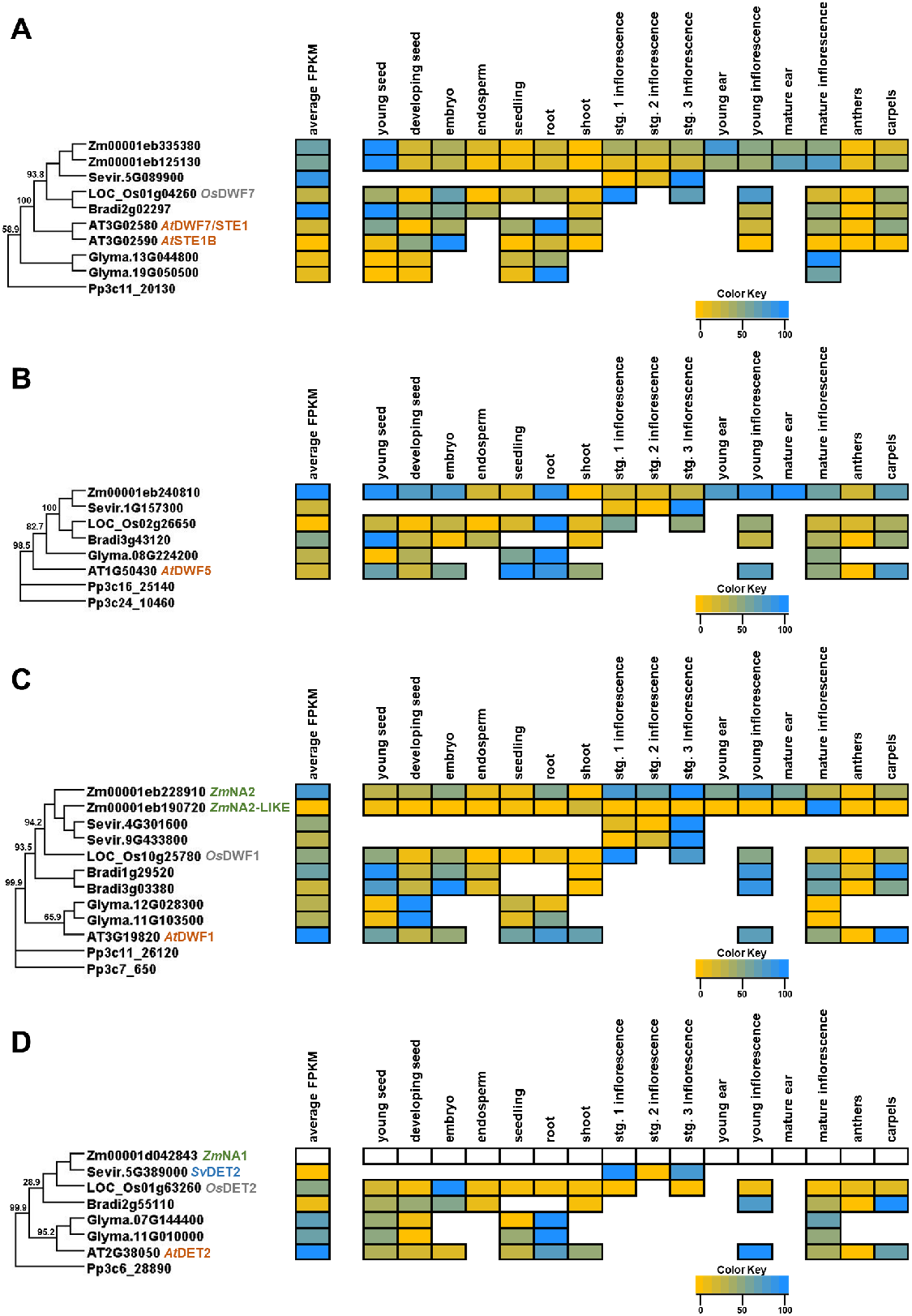
Phylogeny and transcript abundance across different tissues of DWARF7 (DWF7), DWF5, DWF1, and DETIOLATED2 (DET2) phytosterol and brassinosteroid biosynthetic enzyme families. Maximum-approximate-likelihood phylogenetic trees of (A) DWF7, (B) DWF5, (C) DWF1, and (D) DET2 amino acid sequences from maize (Zm), Setaria (Sevir), rice (Os), Arabidopsis (AT), soybean (Glyma), Brachypodium (Bradi), and Physcomitrella (Pp) represented as the outgroup. The relative TPM values from re-analyzed publicly available datasets from different developmental tissues are represented as a heatmap depicted next to each respective gene within the family. The yellow color indicates the lowest individual transcript abundance in each tissue and the dodgerblue color indicates the highest abundance in a given tissue. To compare transcript abundance across genes and species within the family, the average FPKM was determined across all tissues analyzed and is presented in the average FPKM column (left) with the same color distribution as previously described. See Supplemental Tables 1-4 for description of tissue types. The expression boxes for *Zm*NA1 were left blank because there was not a gene annotation in the current maize genome version and therefore the previous version gene annotation was used for phylogenetic analysis.

*DWF7* consists of duplicate pairs in Arabidopsis, soybean, and maize and only a single gene in Brachypodium, rice, and Setaria (Figure 5A). The maize duplicate genes (Zm00001eb335380 and Zm00001eb125130) are expressed across all tissues, with the highest levels in young seeds. Arabidopsis *AtDWF7*/*AtSTE1* and *AtSTE1B* are highly expressed in roots and embryos, respectively, thus suggesting that these genes may have tissue-specific functions. Gene expression disparities between species could be evidence of functional diversification between dicots and monocots, but functional evidence is available only for Arabidopsis which indicated partial functional redundancy between *AtDWF7/STE1* and *AtSTE1B* because the single mutants of *AtDWF7/STE1* displayed a weak phenotype (Choe et al., 1999; Cheon et al., 2010).

Comparing species, overlapping expression patterns suggest a conserved function of *DWF5* in roots (Figure 5B). Mutant analysis of *AtDWF5* indicated its critical role in plant growth and development. The *Atdwf5* mutants displayed dwarf stature with compressed internodes and small-dark-green leaves (Choe et al., 2000). The *DWF5* gene has not been characterized in Brachypodium, maize, rice, Soybean, or Setaria. Development of single mutants with visible phenotypes should be relatively easy to obtain given that they are single-copy genes.

The *DWF1* gene family (Figure 5C) consists of species-specific duplicate gene pairs in maize (*ZmNA2* and *ZmNA2-like*), Setaria, Brachypodium, and soybean; and single copies in Arabidopsis (*AtDWF1*) and rice (*OsDWF1/BRD2*). Available expression data suggested at least one copy in each species is expressed relatively broadly while another is restricted, consistent with some subfunctionalization. For example, *ZmNA2*, like its ortholog in rice (*OsDWF1*), showed highest expression in the inflorescences (Figure 5C) while the *ZmNA2-like* transcript was barely expressed except in roots. However, *ZmNA2-like* was hypothesized to be a pseudogene based on frame-shift mutations (Best et al., 2016). The *Atdwf1* and *Atdwf5* mutants exhibited similar BR-deficient phenotypes and growth defects that were corrected with exogenous treatment of castasterone (CS) or BL (Choe et al., 1999; Inoue et al., 2017; Youn et al., 2018). Functional characterization showed that due to defective sterol C-24 reductase, *Osdwf1*/*brd2* and *Zmna2* mutants were both semi-dwarfed with compressed internodes and erect dark-green short leaves (Hong et al., 2005) (Best et al., 2016). The *Osdwf1*/*brd2* mutants also had fewer and deformed panicles, small spikelets, and shorter rachis branches (Hong et al., 2005; Qin et al., 2018), while *Zmna2* mutants produced shorter, fewer, and feminized tassel branches (Best et al., 2016). Thus, *OsDWF1/BRD2* and *ZmNA2* control both vegetative shoot and reproductive inflorescence architecture. *DWF1* has been extensively investigated in Arabidopsis, maize, and rice but not in other species, suggesting urgency of knockout experiments to understand its evolutionary changes.

In summary, DWF1, DWF5, and DWF7 have been implicated in the regulation of various plant growth and development aspects, which include reproductive development and grain yield (Hong et al., 2005; Best et al., 2016), root development (Choe et al., 1999; Youn et al., 2018), stomatal opening (Inoue et al., 2017), and leaf and internode expansion (Choe et al., 2000; Hong et al., 2005; Best et al., 2016). Consistent with the DWF1-DWF5-DWF7 reactions marking the onset of the BR-dependent pathway, and unlike the upstream mutants described previously, the growth defects of *dwf1*, *dwf5*, and *dwf7* mutants were rescued by exogenous treatment with BL (Clouse, 2002).

#### Campesterol to brassinolide

BL is, in most plants, the most biologically active and abundant BR and therefore considered the terminal product of the BR biosynthetic pathway (Choe, 2006; Wei and Li, 2016). However, recent studies have reported disparities between monocots and dicots in bioactivity of BR compounds (Kim et al., 2008). For instance, in dicots (Arabidopsis and tomato), both BL and its immediate precursor CS (Figure 1) are detected in abundance, but BL was the most active end-product of the pathway (Choe, 2006; Wei and Li, 2016). Conversely, the most active and terminal BR in rice is CS (Kim et al., 2008). By inference, CS may be the most active BR in other monocots like Brachypodium, maize, and Setaria, though tests of this hypothesis are not reported in the current literature.

In overview, plants utilize CR to synthesize CS through either of two, interconnected pathways, campestanol-(CN-) independent and the other CN-dependent (Figure 1; (Wei and Li, 2016; Ohnishi, 2018)). In the CN-independent pathway, CR is converted to 22α-hydroxy-campesterol (22-OHCR), 22α-hydroxy-campest-4-en-3-one (22-OH-4-en-3-one), 22α-hydroxy-5α-campestan-en-3-one (22-OH-3-one), and then 3-epi-6-deoxoCT which is converted to 6-deoxo-TY that converges with the late C-6 oxidation route (Figure 1; (Ohnishi et al., 2012; Vriet et al., 2013; Ohnishi, 2018)).The CN-dependent pathway consists of two interconnected parallel routes, the early and late C-6 oxidation pathways (Ohnishi, 2018). In the early C-6 oxidation route, CN is oxidized to 6-oxocampestanol (6-oxoCN), and then 6-oxoCN is converted to CS through cathasterone (CT), teasterone (TE), 3-dehydroteaserone (3DT), and typhasterol (TY) (Figure 1; (Ohnishi et al., 2012; Ohnishi, 2018)). In the late C-6 oxidation pathway, CN is hydroxylated to 6-deoxocathasterone (6-deoxo-CT) and 6-deoxoCT is oxidized to 6-deoxoteasterone (6-deoxoTE), 3-dehydro-6-deoxoteasterone (6-deoxo3DT), 6-deoxotyphasterol (6-deoxo-TY), 6-deoxocastasterone (6-deoxoCS), and finally to CS (Ohnishi et al., 2012; Ohnishi, 2018). Through these steps, all highly conserved in the plant kingdom, the enzymatic conversion of CR to CS involves reduction, hydroxylation, and oxidation reactions catalyzed by one sterol 5α-reductase and several cytochrome P450s (CYPs). The CYP enzymes that catalyze oxidation and hydroxylation reactions in BR biosynthesis include CYP90A1, CYP90B1, CYP90C, CYP90D, CYP85A, and CYP724 (Figure 1; (Ohnishi et al., 2006; Ohnishi, 2018)).

#### Sterol 5α-reductase

The aforementioned sterol 5α-reductase is encoded by *DEETIOLATED2*/*NANA1* (*DET2*/*NA1*) and it catalyzes the C-5 reduction of campest-4-en-3-one (4-en-3-one), 22-OH-4-en-3-one, and 22,23-diOH-4-en-3-one to yield 5α-campestan-3-one (3-one), to 22-OH-3-one, and 6-deoxo3DT, respectively (Ohnishi et al., 2012; Ohnishi, 2018; Wei and Li, 2020). In all species examined here except soybean, DET2/NA1 is encoded by a single gene (Figure 5D). *DET2*/*NA1* has been well-characterized only in Arabidopsis (*AtDET2*) and maize (*ZmNA1*) (Chory et al., 1991; Choe et al., 1999; Hartwig et al., 2011). The *Atdet2* and *Zmna1* mutants exhibited similar growth and developmental defects compared to *Atdwf1* and *Zmna2,* respectively (Hartwig et al., 2011; Noguchi et al., 1999). However, the *Atdet2* and *Zmna1* pleiotropic phenotypes were associated with a defective 5α-reductase that accumulates 4-en-3-one substrates downstream of *At*DWF1/*Zm*NA2 (Noguchi et al., 1999; Hartwig et al., 2011).

#### C-22 hydroxylase

CYP90B/DWF4 and CYP724A/D11 catalyze the C-22 hydroxylation of CR, 4-en-3-one, 3-one, or CN to yield 22-OHCR, 22-OH-4-en-3-one, 22-OH-3-one, or 6-DeoxoCT, respectively (Figure 1 and 6; (Ohnishi, 2018)). Consistent with previous studies (Ohnishi et al., 2006; Ohnishi, 2018), our phylogenetic analysis (Figure 6) indicated that *CYP90B/DWF4* and *CYP724A/D11* assort into two clustered subclades (Ohnishi et al., 2006; Ohnishi, 2018). In the species examined in this review, *CYP90B/DWF4* and *CYP724A*/*D11* are single copy genes except for *CYP90B/DWF4* with three copies in soybean (Figure.6). CYP90B/DWF4 and CYP724A/D11 both catalyze the C-22 hydroxylation of BRs. For instance, Arabidopsis *Atdwf4* mutants exhibited growth defects indistinguishable from *Atdet2* or *Atdwf1* (Choe et al., 1998). However, overexpression of D11 (*At*CYP724A1) complemented *Atdwf4* mutants (Choe et al., 1998; Zhang et al., 2012). In rice, both *Osdwf4* and Os*d11* loss of function mutants displayed mild phenotypic defects, such as semi-dwarfed stature with erect leaves and small-round grains. *Osdwf4*; *Osd11* double mutants were extremely dwarfed with deformed and erect leaves, suggesting that OsDWF4 and OsD11 function somewhat redundantly (Sakamoto et al., 2006). In Setaria, the *CYP724A/D11* homolog *Svbsl1* loss of function mutants were primarily affected in the inflorescence and were semi-dwarf with shorter and fewer panicles and bristles. This was associated with failed bristle initiation and spikelet meristem determinacy and maintenance (Yang et al., 2018). Mutants of *ZmD11* have not been reported but gene function has been tested in other species; whereas overexpression of *ZmD11* and *ZmDWF4* restored the growth of *Osd11* and *Atdwf4* mutants, respectively, thus suggesting functionality of DWF4/CYP90B and CYP724A/D11 is conserved across plant species (Liu et al., 2007; Sun et al., 2021). Overexpression of *Zm*D11 in rice also affected grain size and quality (Sun et al., 2021). In our data mining analysis *DWF4/CYP90B* and *CYP724A/D11* transcripts were expressed highly in different tissues, specifically vegetative, compared to reproductive tissues (Yang et al., 2018), despite their overlapping catalytic functionality. For instance, *ZmD11*, *OsD11*, and *SvBSL1* were highly expressed in young ears, panicles, and developing inflorescence primordia, respectively (Wu et al., 2016; Yang et al., 2018; Sun et al., 2021). The *ZmDWF4*, *OsDWF4*, and *Sevir.9G483600* genes were highly expressed in young shoots, leaf blades, and leaves, respectively (Sakamoto et al., 2006; Liu et al., 2007; Yang et al., 2018; Sun et al., 2021).

**Figure 6.**
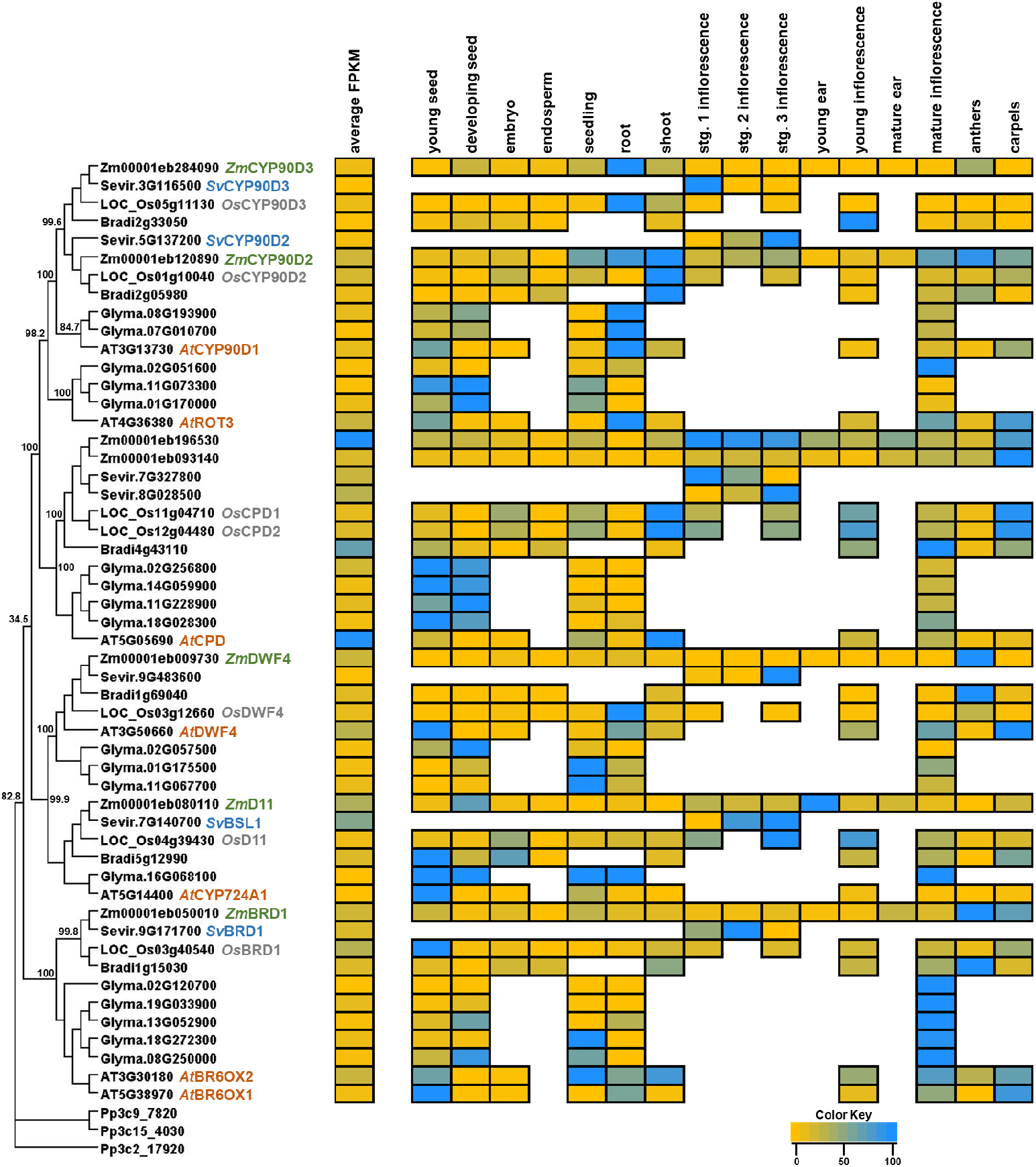
Phylogeny and transcript abundance across different tissues of the CYTOCHROME P450 (CYP450) enzyme family involved in multiple steps of brassinosteroid biosynthesis. Maximum-approximate-likelihood phylogenetic tree of brassinosteroid biosynthetic CYP450 amino acid sequences from maize (Zm), Setaria (Sevir), rice (Os), Arabidopsis (AT), soybean (Glyma), Brachypodium (Bradi), and Physcomitrella (Pp) represented as the outgroup. The relative TPM values from re-analyzed publicly available datasets from different developmental tissues are represented as a heatmap depicted next to each respective gene within the family. The yellow color indicates the lowest individual transcript abundance in each tissue and the dodgerblue color indicates the highest abundance in a given tissue. To compare transcript abundance across genes and species within the family, the average FPKM was determined across all tissues analyzed and is presented in the average FPKM column (left) with the same color distribution as previously described. See Supplemental Tables 1-4 for description of tissue types.

#### C-23 hydroxylase

The C-23 hydroxylation of 22-hydroxylated BRs to yield 23-hydroxylated BRs is controlled by ROTUNDIFOLIA3 (ROT3)/CYP90C and CYP90Ds that function redundantly as C-23 hydroxylases (Ohnishi et al., 2006). ROT3/CYP90C and CYP90Ds predominantly convert 6-deoxoTE to 3-dehydro-6-deoxoTE, TE to 3-dehydroTE, 3-epi-6-deoxoCT to 6-deoxoTY, 22-OH-4-en-3-one to 3-dehydro-6-deoxoTE, 6-deoxoCT to 6-deoxoTE, and 6-deoxoTE to 3-dehydro-6-deoxoTE (Sakamoto et al., 2012; Ohnishi, 2018). Consistent with previous studies (Rizal et al., 2015), our phylogenetic analysis (Figure 6) indicated that ROT3/CYP90C1 and CYP90D1 are restricted to dicots (Arabidopsis and soybean), while CYP90D2 and CYP90D3 are specific to monocots (maize, Setaria, Brachypodium, and rice) due to a duplication event that occurred post-divergence between monocots and dicots and before the evolution of the grasses. In Arabidopsis, CYP90C and CYP90D enzymes are encoded by *ROT3*/*CYP90C1* and *CYP90D1*, respectively (Kim et al., 2005; Ohnishi et al., 2006). Mutant studies indicate that *ROT3*/*CYP90C1* and *CYP90D1* encode functionally redundant C-23 hydroxylases. *Atrot3* mutants displayed weak BR-deficient phenotypes, and *Atcyp90d1* mutants appeared similar to wild-type plants but the *Atrot3;cyp90d1* double mutants displayed extremely dwarf phenotypes indistinguishable from *Atdet2* and *Atdwf4* mutants (Kim et al., 2005; Ohnishi et al., 2006). Gene expression data further support the partial functional redundancy of ROT3/CYP90C1 and CYP90D1 both are expressed in most tissues. The *At*ROT3 transcripts accumulate to slightly higher levels than *AtCYP90D1*. In monocots, the C-23 hydroxylation reactions are catalyzed by CYP90D2 and CYP90D3 enzymes, encoded by a duplicate gene pair (Figure 6; (Hong et al., 2003; Sakamoto et al., 2012; Rizal et al., 2015)). The CYP90D2 and CYP90D3 proteins have overlapping functions; for instance, CYP90D2 loss of function mutants in rice and sorghum have weak growth defects compared to other BR-deficient mutants (Hong et al., 2003; Sakamoto et al., 2012; Rizal et al., 2015), which could be due to partial complementation by CYP90D3. Similarly, the single mutants of maize genes that are orthologs of *OsCYP90D2* and *OsCYP90D3* exhibit weak phenotypes with no obvious BR-related growth defects, but double mutants are extremely dwarfed and indistinguishable from *Zmna1* and *Zmna2* mutants (B. Zebosi and E. Vollbrecht, unpublished). These data further affirm that *Zm*CYP90D2 and *Zm*CYP90D3 function redundantly as C-23 hydroxylases. Unequal functional redundancy between *ZmCYP90D2* and *ZmCYP90D3* is supported by expression data (Figure 6) wherein *ZmCYP90D2* was more predominantly expressed than *ZmCYP90D3* across tissues, except in roots.

#### C-3 oxidases

CYP90As function as C-3 oxidases and mediate the C-3 oxidation of CN, 22-OHCR, and (22R,23R)-22,23-Dihydroxycampesterol to produce 4-en-3-one, 22-OH-4-en-3-one, (22R,23R)-22,23-dihydroxy-campest-4-en-3-one, respectively (Ohnishi et al., 2012). CYP90A in Arabidopsis is encoded by *CONSTITUTIVE PHOTOMORPHOGENIC DWARFISM* (*CPD*) (Ohnishi et al., 2012) and *CPD* gene copy numbers vary (Figure 6). Arabidopsis and Brachypodium have a single CPD, soybean have four, while monocots (maize, Setaria, and rice) have two CPDs as a result of gene duplication (Figure 6). Gene expression analysis (Figure 6) across the *CPD*/*CYP90A* subclade shows that *AtCPD* has the highest average expression compared to the orthologs in the other species. Nevertheless, the *CPD* genes across species exhibit expression pattern similarities. Transcripts of *At*CPD and its co-orthologs in rice (*Os*CPD1 and *Os*CPD2) are highly expressed in shoot tissues and carpels. The rice *OsCPD* genes (*OsCPD1* and *OsCPD2*) (Zhan et al., 2022) and their maize orthologs are similarly expressed in carpels, suggesting some functional conservation between maize and rice. Soybean CPDs are highly and overlappingly expressed in young and developing seeds. Within species, especially maize, rice, soybean, and Setaria, the duplicate *CPD* genes display overlapping tissue-expression patterns, thus suggesting functional redundancy. In Arabidopsis, *AtCPD* has been extensively characterized through mutant screens and metabolite profiling, and the *Atcpd* mutants exhibited extreme abnormal growth phenotypes similar to those reported for *Atdwf1* and *Atdet2* mutants (Szekeres et al., 1996; Ohnishi et al., 2012). In addition, the *Atcpd* mutants accumulated 22-OHCR and *Atcpd*; *Atdet2* double mutants indicated that *At*CPD functions early in the BR pathway and upstream of *At*DET2 (Ohnishi et al., 2012). Recently, *OsCPD* loss of function mutants (*Oscpd1* and *Oscpd2*) were characterized in rice; neither single mutant have obvious growth defects, but the double mutants (*oscpd1*;*oscpd2*) display multiple developmental defects similar to *Osdwf4*; *Osd11* double mutants (Zhan et al., 2022), thus implying functional redundancy. To date, functional evidence of CPD genes in maize, soybean, Brachypodium ,and Setaria remain limited due to a lack of mutants. Further research is needed to develop mutants of these genes, and double mutant combinations will need to be developed due to the likely redundancy of gene function within species.

#### C-6 oxidases

CYP85A/BR-C-6 oxidase catalyzes the oxidation of CS to BL. CYP85A also mediates several intermediate reactions and these include the enzymatic conversion of 6-deoxoTE, 6-deoxo3DT, 6-deoxo-TY, and 6-deoxoCS to TE, 3DT, TY, and CS (Figure 1).*CYP85A*/*BR6OX* genes encode BR-C-6 oxidase, and their copy number varies between dicots and monocots. Monocots have a single gene copy (dubbed CYP85A1), whereas Arabidopsis contain two gene copies (CYP85A1 and CYP85A2) and soybean has five CYP85As (Jager et al., 2007; Kim et al., 2008). However, expression data shows that *CYP85A* genes exhibit overlapping and organ-specific expression patterns. When averaged over tissues, *CYP85A*s have the highest expression in young seeds except for *ZmBRD1* and Bradi1g1503, which are expressed strongly in the anthers, suggesting functional diversification (Figure 6). Unlike in Setaria, soybean, and Brachypodium, CYP85A loss of function mutants have been well characterized in Arabidopsis, maize, and rice.

Mutant studies and metabolic analyses in Arabidopsis indicated that *At*CYP85A1/*At*BR6OX1 and *At*CYP85A2/*At*BR6OX2 catalyze similar reactions yielding BL, except the conversion of CS to BL is uniquely presided over by *At*BR6OX2 (Kim et al., 2005). Comparing enzymatic activity between these enzymes, *At*BR6OX2 had a measurably higher oxidase activity than *At*BR6OX1, consistent with previous mutant studies (Kim et al., 2005). For instance, *Atbr6ox1* mutants have no growth defects, while *Atbr6ox2* mutants had mild defects similar to those of other BR-deficient mutants (Kim et al., 2005). Growth defects were amplified, however, in the *Atbr6ox1*;*Atbr6ox2* double mutants (Kim et al., 2005; Kwon et al., 2005). In addition, *Atbr6ox2* and *Atbr6ox1*;*Atbr6ox2* mutant defects were entirely restored with exogenous application of BL and CS, emphasizing that BL and/or CS activity is required for Arabidopsis growth and development. Mutant phenotypic differences between *Atbr6ox1* and *Atbr6ox2* could suggest that the CS levels are supplemented by *At*BR6OX1 in the *Atbr6ox2* mutants; however, the levels are insufficient to maintain proper plant growth and development (Kim et al., 2005). On the other hand, the phenotypic differences could imply that CS and BL function overlappingly but with some tissue specificity; for instance, BL was found to be enriched in reproductive tissues (flowers, seeds, and siliques), while CS accumulated more in vegetative tissues (roots, shoots, and leaves) (Shimada et al., 2003; Kim et al., 2005; Nomura et al., 2005). The *At*BR6OX1 and *At*BR6OX2 also display overlapping and tissue-specific expression patterns (Figure 6); for instance, *At*BR6OX2 is highly expressed in most tissues except in the young seeds and carpels, emphasizing the mutant phenotypic disparities and tissue-specific functionalization between *AtBR6OX1* and *AtBR6OX2*.

Loss of function of *ZmBRD1* and *OsBRD1* in maize and rice, respectively, showed various pleiotropic vegetative and reproductive defects, highlighting the essentiality of CYP85A1 in BR biosynthesis and plant development (Mori et al., 2002; Makarevitch et al., 2012). *Zmbrd1* mutants were extremely dwarfed with inflorescence defects and feminized tassels indistinguishable from *Zmna1* and *Zmna2* mutant phenotypic defects (Makarevitch et al., 2012), underlining that BRs modulate sex determination and reproductive architecture. *ZmBRD1* also regulates organ boundary formation, cell division, and expansion as mutants had enlarged auricles and less distinct leaf auricle-blade boundary, disorganized cell files, and reduced cell expansion (Baluška et al., 2001; Makarevitch et al., 2012; Castorina et al., 2018). These mutants also exhibited BR-deficient-related defects at the seedling level and were partially rescued with the exogenous treatment of BL (Makarevitch et al., 2012). Plant growth and developmental defects of *Osbrd1* mutants were similar to those reported for *Zmbrd1* mutants (Hong et al., 2002; Makarevitch et al., 2012), indicating plausible functional conservation of *BRD1* gene function between maize and rice. Despite partially rescuing *Zmbrd1* and *Osbrd1* phenotypic defects by exogenous BL treatment, recent studies have shown that monocots likely use primarily CS instead of BL as the bioactive BR. CS was shown to be the end-product of BR biosynthesis in rice, and it was noted that rice (and all monocots) lacks a CYP85A2 that may more specifically catalyze the conversion of CS to BL (Kim et al., 2008; Roh et al., 2020). To determine more universally whether CS or BL is the end-product of BR biosynthesis in a range of monocots, there is a need to carefully profile metabolite accumulation levels and quantify the extent of growth rescue by applying BL and CS in CYP85A loss of function mutants.

### Brassinosteroid Movement

BRs are synthesized within the cell, primarily at the endoplasmic reticulum (Klahre et al., 1998; Shimada et al., 2001), and then perceived at the cell membrane surface outside of the cell. Therefore, local transport is necessary for bio-active BRs to exit the cell and bind to the BR receptor anchored in the cell membrane. It is not yet known how BRs move out of the cell or even through the intracellular environment from the ER membrane to the plasma membrane for export. It has been hypothesized that either movement could occur by creating BR conjugates or binding of BR to protein complexes to reduce the hydrophobic state of BRs and facilitate their movement (Fujioka and Yokota, 2003; Marković-Housley et al., 2003; Sasse, 2003; Choe, 2006). Further studies are necessary to elucidate how intracellular BRs reach the cell membrane after their synthesis. Once BRs reach the cell membrane, to exit the cell they need to cross the lipid bilayer by an either active, passive, or combined mechanism. Symplastic transport of BR precursors cell-to-cell via plasmodesmata has now been demonstrated (Nolan et al., 2020) but export of bioactive BRs to the apoplast is necessary to initiate the BR signaling cascade (Vukasinovic and Russinova, 2018). To date, there have been no active transporters of BRs across the cell membrane identified in any plant species. ATP-binding cassette (ABC) transporters have been shown to actively transport steroidal compounds out of animal cells (Agassandian et al., 2004; Tarling and Edwards, 2011), so this should be a topic for future research. Given that BRs can feedback-regulate biosynthesis and signaling components (Wei and Li, 2020), BRs can likely bind to receptors of the same cell that synthesized them. Moreover, once outside of the cell BRs could also undergo local short-distance transport to affect nearby cells. Apoplastic movement of BRs should also be possible for BRs to influence nearby cells within a given tissue or organ.

Long-distance transport of BRs has been studied in a few species, including tomato and pea (Symons et al., 2008). Exogenous application of BRs to the roots can recover biosynthetic mutant phenotypes evident in the shoot in Arabidopsis (Choe et al., 1998). However, grafting experiments in both tomato and pea showed that grafting wild-type rootstocks to mutant scions could not recover the dwarf shoot phenotype, thus indicating that *in vivo* synthesized BRs were not acropetally transported (Symons and Reid, 2004; Montoya et al., 2005). Furthermore, grafting wild-type scions to mutant root stocks did not increase the endogenous levels of BRs in the root stock, indicating a lack of long distance basipetal transport (Symons and Reid, 2004; Montoya et al., 2005). Tissue excision experiments also showed that removing a single organ had no effect on the endogenous levels of BRs in another organ (Symons and Reid, 2004). Taken together, there is no evidence that endogenous BRs are transported long-distances throughout a plant. There has not been extensive research in Arabidopsis, maize, rice, Setaria, Brachypodium, or soybean on long-distance transport of BRs. Further research is needed to understand BRs transport throughout plants during growth and development.

### Brassinosteroid Catabolism and Homeostasis

BR homeostasis is comprised of a complex interplay among spatiotemporal distribution of active BR compound(s), perception of, signaling and response to those BRs, and inputs from genotype and environment. Given that no mechanism of long-distance active transport of BRs has been described (Symons et al., 2008), local metabolism largely determines tissue composition of BRs; this BR metabolism includes biosynthesis, as discussed, as well as catabolic processes like conjugation, modification, and degradation. Catabolism in vivo of BL and CS was first documented by the discovery in Arabidopsis of the C26 hydroxylase PHYB ACTIVATION-TAGGED SUPPRESSOR1 (BAS1) which when overexpressed, mimicked BR biosynthetic and response mutants and caused reduced levels of BRs (Neff et al., 1999). BAS1 is single copy in Arabidopsis, maize, Setaria, and Brachypodium and there are three copies in soybean (Supplemental Figure 1). The highest average expression across species is in developing and mature inflorescences, but Arabidopsis and one soybean transcript (Glyma.06G215200) also have high expression in seedlings.

Genetically related C26 hydroxylases have since been analyzed in many plant species. Several different conjugating and modifying activities, that act either directly on the predominant bioactive compounds BL and CS and/or on biosynthetic intermediates, have now been documented including glycosyltransferases, a reductase, acyltransferases, and sulfotransferases (Vriet et al., 2013; Wei and Li, 2020). In addition to biochemical evidence, *in planta* genetic evidence has demonstrated for many catabolic genes, overexpression mimics BR-deficient phenotypes, while loss-of-function mutants show organ elongation or other BR responsive phenotypes (Neff et al., 1999; Turk et al., 2005; Thornton et al., 2010; Schneider et al., 2012; Zhuang et al., 2022). Mechanisms that regulate BR catabolic genes during plant development and in response to external cues have also begun to be elucidated (Wei and Li, 2020).

### Brassinosteroid Perception

#### BRASSINOSTEROID INSENSTIVE1/BRASSINOSTEROID INSENSITIVE LIKE (BRI1/BRL)

BRs are perceived at the plasma membrane through binding of bio-active BRs by the BRASSINOSTEROID INSENSITIVE1 (BRI1) and several BRASSINOSTEROID INSENSITIVE1-LIKE (BRL) receptor kinases (Clouse et al., 1996; Li and Chory, 1997; Cano-Delgado et al., 2004) (Figure 7). Within the BRI1 and BRL proteins is an extracellular domain that is necessary to bind BRs and consists of multiple leucine rich repeat (LRR) motifs and a signature island domain (ID) positioned near the carboxy terminus of the LRR domain (Kinoshita et al., 2005; Hothorn et al., 2011; She et al., 2011). These proteins also contain a transmembrane sequence, responsible for anchoring in the plasma membrane, as well as a protein kinase domain positioned in the cytoplasm that triggers intracellular signal responses.

**Figure 7.**
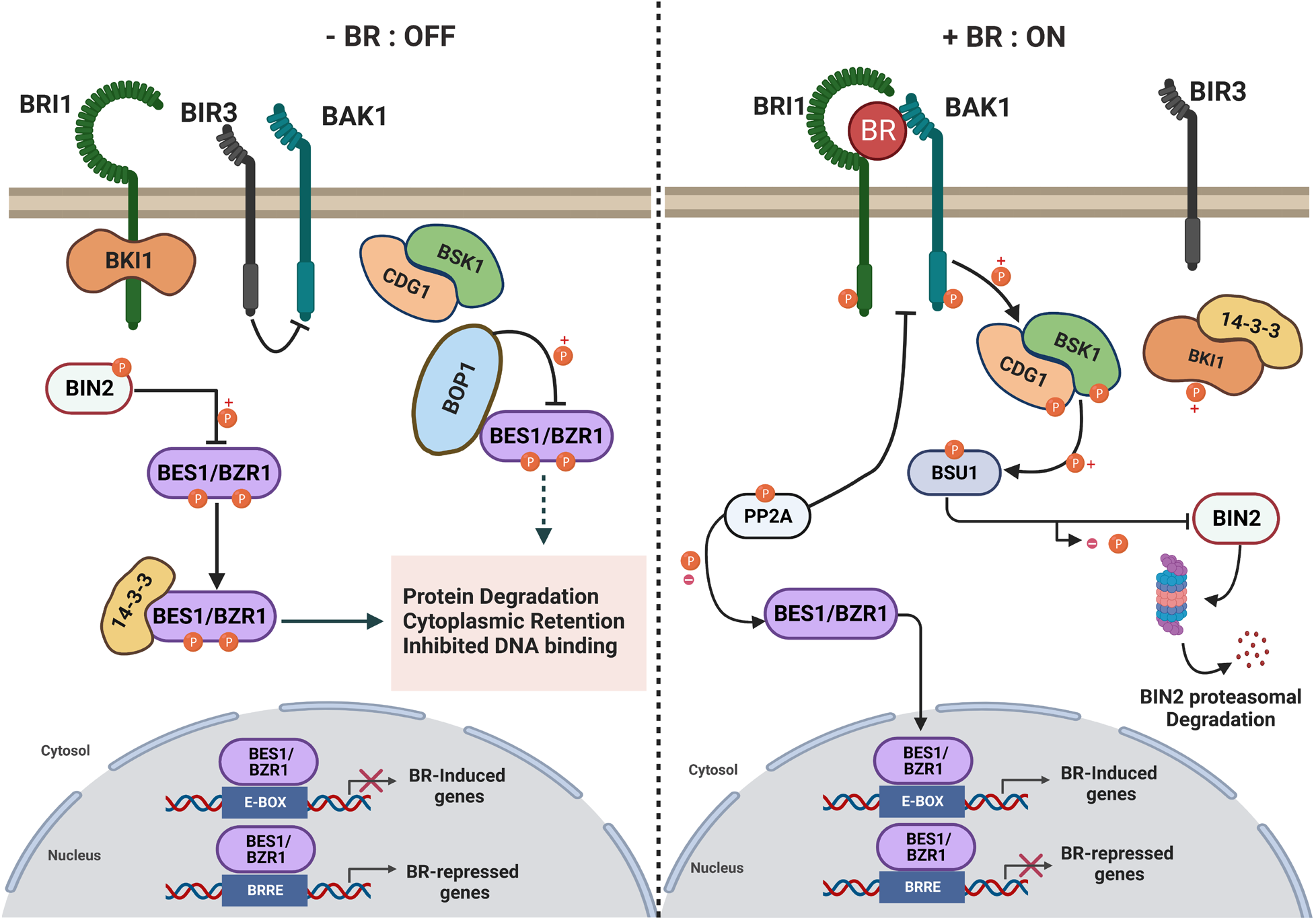
Brassinosteroid signaling pathway. The brassinosteroid (BR) signaling pathway shown when BRs are not present on the left and when present on the right. This figure was modified according to (Nolan et al., 2020) and created using biorender software.

There are four members of the BRI1/BRL family in Arabidopsis, rice, Brachypodium, and Setaria; five members in maize; and six in soybean. (Figure 8A). Most duplications and divergences in this family predate the split between monocots and eudicots, except for *BRI1* homologs. *BRI1* orthologs are single copy in rice and Setaria but recent genome duplications in maize, Brachypodium, and soybean led to one additional *BRI1* gene via duplication of the closest homolog to *AtBRI1* followed by retention of both copies. The second Brachypodium *BRI1* gene is likely a pseudogene as it encodes variant amino acids at several, otherwise extremely well conserved residues in the kinase domain. Across species, *BRI1* genes are more highly expressed than *BRL* gene family member(s), with the highest BRI1 expression in developing inflorescences and floral tissues. *BRL* genes are most highly expressed in root tissue, except for *Os*BRL2 which is highest in young inflorescences. These similar *BRI1*/*BRL* expression patterns similar across species suggest an ancient, conserved sub-functionalization among monocots and eudicots.

**Figure 8.**
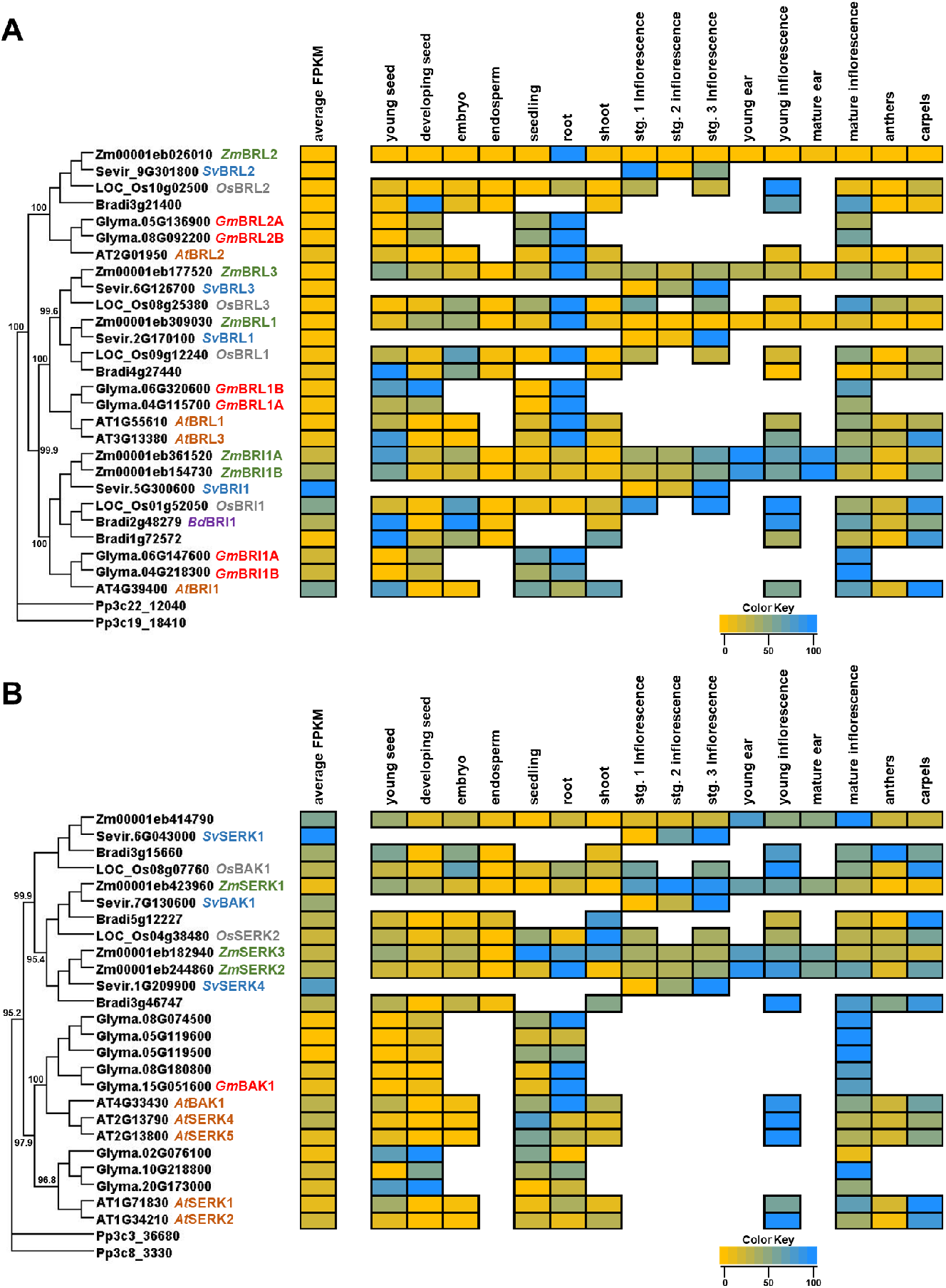
Phylogeny and transcript abundance across different tissues of BRASSINOSTEROID INSENSITIVE1 (BRI1) and BRI1-ASSOCIATED RECEPTOR KINASE families involved in brassinosteroid perception. Maximum-approximate-likelihood phylogenetic trees of (A) BRI1 and (B) BAK amino acid sequences from maize (Zm), Setaria (Sevir), rice (Os), Arabidopsis (AT), soybean (Glyma), Brachypodium (Bradi), and Physcomitrella (Pp) represented as the outgroup. The relative TPM values from re-analyzed publicly available datasets from different developmental tissues are represented as a heatmap depicted next to each respective gene within the family. The yellow color indicates the lowest individual transcript abundance in each tissue and the dodgerblue color indicates the highest abundance in a given tissue. To compare transcript abundance across genes and species within the family, the average FPKM was determined across all tissues analyzed and is presented in the average FPKM column (left) with the same color distribution as previously described. See Supplemental Tables 1-4 for description of tissue types.

*Atbri1* mutants display a severe dwarfism phenotype. The first *Atbri1* mutants were discovered in mutant screens to identify loci that were insensitive to epi-BL treatment (Clouse et al., 1996; Li and Chory, 1997). In addition to extracellular LRR and ID domains essential to bind and perceive BRs at the cell membrane (Kinoshita et al., 2005; She et al., 2011), BRI1 proteins contain a signal peptide, a putative leucine-zipper motif, a transmembrane domain, and a cytoplasmic kinase domain that all contribute to BRI function in perceiving BRs and initializing the phosphorylation cascade necessary for BR signaling (Li and Chory, 1997). The BRL gene products, *At*BRL1 and *At*BRL3, were also shown to bind BL with high affinity. Overexpression lines of *At*BRL2 were not able to bind BL. *At*BRL1 and *At*BRL3 are partially redundant with *At*BRI1 in regulating growth and development of Arabidopsis plants (Cano-Delgado et al., 2004; Zhou et al., 2004). Transgenic expression of *Gm*BRI1A in the *Atbri1-5* mutant background (Wang et al., 2014) and overexpression of *Gm*BRI1B in the *Atbri1-6* mutant background (Peng et al., 2016) was able to rescue the Arabidopsis mutants’ phenotype, indicating a conservation of function across species.

To initially characterize the BRI1/BRL gene family in maize, RNA-interference (RNAi) transgenic plants created to knock-down transcript levels (Kir et al., 2015). The RNAi constructs were designed to primarily target *ZmBRI1A* and *ZmBRI1B*, but transcript levels were reduced for all five *BRI1*/*BRL* homologs. The BRI1-RNAi lines also had increased expression of the *ZmBRD1* and *AtCPD-like* (Zm00001eb196530) transcripts and thus likely accumulated bio-active BRs to increased levels, although levels were not analyzed. The *Zm*BRI1-RNAi lines were moderately dwarfed due to inhibited elongation of all internodes, especially for upper internodes between the ear and tassel. Leaf blade and leaf sheath length was also affected, but leaf blade width was not (Kir et al., 2015). Interestingly, despite knockdown of all *BRI1*/*BRL* homologs, none of the transgenic maize lines recovered by the transgenic RNAi approach showed the extreme dwarfism of *bri1* single mutants from Arabidopsis or rice, suggesting a need for mutant analysis in maize.

The *OsD61* gene encodes for *Os*BRI1 in rice and the *Osd61* mutant phenotype closely resembles the biosynthetic mutant *Osbrd1-1* (Nakamura et al., 2006). The severe *Osd61-4* allele has a mutation in the kinase domain of *Os*BRI1 and demonstrates *OsD61* is necessary to regulate BR signaling (Zhao et al., 2013). *Osd61* mutant plants were severely dwarfed with abnormal leaves and were sterile due to the absence of flowers. RT-PCR analysis showed that *OsBRI1* was primarily expressed in shoots while *OsBRL1* and *OsBRL3* were expressed in roots, as we observed in our expression analysis (Figure 8A). CS levels were close to 30 times higher in *Osd61* mutant shoot tissue as compared to wild-type siblings, but only 1.5 times higher in root tissue. The reduction in mutant root length was also much milder compared to *Osbrd1-1* (Nakamura et al., 2006). These results suggest that in root tissue other receptors like *Os*BRL1 and *Os*BRL3 may be sufficient or may compensate for the lack of *Osd61* function.

Transgenic RNAi lines have also been developed that target and lower expression of *BdBRI1* transcripts (Feng et al., 2015). The RNAi lines exhibited stunted growth but not extreme dwarfism, like what was observed in maize. The *BdBRI1* RNAi lines were also more drought tolerant and exhibited altered expression of drought-responsive genes under drought conditions (Feng et al., 2015) highlighting a potential applied usage of BR mutants for drought tolerance across species. Surprisingly, overexpression of *BdBRI1* in an *Atbri1-5* background did not recover the mutant phenotype (Corvalán and Choe, 2017). The function of BRI1/BRL proteins have not been studied in Setaria and future work is needed to better elucidate the possible diversification or conservation of these genes in BR signaling.

#### BRI1-ASSOCIATED RECEPTOR KINASE (BAK1)

The *BAK1* gene encodes an LRR receptor-like kinase similar to BRI1. The BAK1 kinase is in the somatic embryogenesis receptor kinase (SERK) family of LRR kinases. BAK kinases contain a signal peptide that targets the plasma membrane, an extracellular LRR domain, a transmembrane domain, and a cytosolic receptor like kinase domain (Li et al., 2002). Four other Arabidopsis genes show high sequence similarity to *AtBAK1* and all five genes are highly expressed in developing inflorescences, but *AtBAK1* is also highly expressed in seedling roots (Figure 8B). The four *BAK1*-like genes in maize also show high expression in developing inflorescences. Phylogenetic analysis suggests that in contrast to the ancient conservation of *BRI1* genes, the *BAK1* gene family underwent independent expansion in eudicots and in grass species (Figure 8B). The *ZmSERK2* and *ZmSERK3* genes, which show an expression pattern like that of *AtBAK1*, also have high expression in seedling roots possibly indicating a redundancy of function in maize. There are eight *BAK1*-like genes in soybean. In rice two genes that encoded polypeptides with sequence similarity to *AtBAK1*/*SERKs*. Three additional genes (LOC_Os08g07890, LOC_Os06g12120 and LOC_Os08g07890) show phylogenetic and gene structure features suggesting altered functions (Man et al., 2020) and were left out of our phylogenetic and expression analyses. *OsBAK1* gene expression is higher in developing inflorescences and *OsSERK2* is higher expressed in seedling tissue, suggesting some functional diversification in regulating BR signaling. Setaria only has three *BAK1*-like genes and *SvSERK1* and *SvSERK4* have higher average expression than *SvBAK1*, in the few tested tissues. The three *BAK1*-like genes in Brachypodium show a similar expression profile to genes within their respective subclades.

The *AtBAK1* gene was discovered by screening for suppressors of the *Atbri1* mutant. Dominant *Atbak1-1D* mutants suppressed the *Atbri1-5* short inflorescence, shortened petiole and curled lamina phenotypes. *Atbak1-1D* single mutants had longer hypocotyls, stems, and rosette width as compared to wild-type siblings. The loss of function mutant, *Atbak1-1*, was semi-dwarfed with reduced sensitivity to BL-induced root length inhibition. BAK1 localizes to the plasma membrane and physically interacts with BRI1 *in vitro* and *in vivo*. When BRs are not present, BRI1 and BAK1 are not phosphorylated and do not interact. When BRs bind to BRI1 it associates with BAK1 and both become auto- and/or trans-phosphorylated, kicking off an intracellular phosphorylation cascade to activate BR signaling and response (Li et al., 2002). *At*SERK1 is functionally redundant with *At*BAK1 in BR signaling. *At*BAK1 also acts as a co-receptor to regulate plant innate immunity (Albrecht et al., 2008); however, some immune responses are regulated by BRs independent of *At*BAK1 (Albrecht et al., 2012; Belkhadir et al., 2012).

Loss of function *Atbak1* mutants have short primary roots and making double or triple mutants with loss of function *Atserk1* and/or *Atserk2* alleles further decreased the primary root length (Du et al., 2012; Ou et al., 2022). Root growth and development has been shown to be regulated by *At*ROOT MERISTEM GROWTH FACTOR1s (RGFs)/GLOVEN(GLV)/CLE-like (CLEL) (Fernandez et al., 2013), a class of small peptide hormones, that are perceived by leucine-rich repeat receptor-like kinases, *At*RGF1 INSENSITIVEs (RGIs) (Ou et al., 2016; Shinohara et al., 2016; Song et al., 2016). These RGIs act as co-receptors with *At*BAK1 and other *At*SERKs to bind RGFs and regulate the root meristem cell niche, lateral root development, and root gravitropism (Ou et al., 2022). The RGF/RGI regulation of root growth has not been directly linked to BRs thereby suggesting *At*BAK1/SERK proteins regulate root growth and development by both BR-dependent and -independent pathways.

Arabidopsis BAK1-INTERACTING RECEPTOR-LIKE KINASE2 (BIR2) and BIR3 gene products were also shown to interact with *At*BAK1 and *At*BRI1 *in vivo* to regulate their function (Halter et al., 2014; Imkampe et al., 2017). *At*BIR3 stabilizes *At*BAK1 and prevents it from associating with ligand binding receptors to form a receptor complex (Imkampe et al., 2017). *At*BAK1, along with *At*SERK4, was also shown to mediate programmed cell death independent of BRs. Also independent of BRs, *At*SERK1 and *At*SERK2 were responsible for male microsporogenesis. A function and/or phenotypic consequence of *At*SERK5 has yet to be described (Albrecht et al., 2008). In soybean, *Gm*BAK1 was shown to form a complex with *Gm*FLAGELLIN-SENSITIVE2 to perceive flagella from *Ralstonia solanacearum*, which also appears to be independent of BRs (Wei et al., 2020).

Maize *BAK1*/*SERK* genes have not yet been characterized through mutant analyses. *Zm*SERK1 and *Zm*SERK2 transcript abundance published in initial studies (Baudino et al., 2001) matches available transcriptomics data (Figure 8B). Transcriptomic studies on somatic embryogenesis in maize identified Zm00001eb414790 as being up-regulated, but neither *Zm*SERK1, *Zm*SERK2, nor *Zm*SERK3 were differentially regulated during this process (Ding et al., 2020). This surprising result may suggest some level of diversification between Arabidopsis and maize for the function of *BAK1*/*SERK* genes (Salaj et al., 2008).

To identify the function of *BAK1*/*SERK* genes in rice growth and development, RNAi knockdown lines were developed. Multiple constructs targeted *OsBAK1*, *OsSERK2*, and the additional SERK-like gene *OsORK1* (LOC_Os08g07890). Variable effects were observed across RNAi events. Severe phenotype events with no detectable expression of *OsBAK1* were dwarfed and insensitive to BL-induced laminar leaf bending. Weak events did have expression of *OsBAK1* but were knocked down in *OsORK1*. These lines were sensitive to BL treatment for laminar leaf bending but lacked strong morphological effects on growth and development (Park et al., 2011). Overexpression in Arabidopsis of *OsBAK1* and *OsSERK2*, but not of *OsSERK3* (LOC_Os06g12120) or *OsSERK4* (LOC_Os02g18320), complemented mutant phenotypes of a weak *Atbri1* allele indicating conservation of *BAK1*/*SERK1* molecular function across species (Li et al., 2009), despite their phylogenetic history (Figure 8B). The Osserk2 mutant has been cloned and showed altered response to BL treatment (Dong et al., 2020). Thus, *OsBAK1* and *OsSERK2* genes are most closely related to *AtBAK1* and are redundant in regulating BR signaling. Future work is needed to investigate the function of *BAK1*/*SERK* genes in Setaria, Brachypodium, and soybean.

#### BRI1 KINASE INHIBITOR1 (BKI1)

The BKI1 protein inhibits BRI1 function when BRs are not present to ensure BRI1 specifically regulates BR signaling (Wang and Chory, 2006; Wang et al., 2014). The *BKI1* gene is part of the larger MEMBRANE-ASSOCIATED KINASE REGULATOR (MAKR) protein family whose members have been implicated in root growth development and responses and whose expression is regulated by multiple phytohormones (Novikova et al., 2021). *BKI1* encodes two highly conserved motifs that regulate its function, the BRI1-interacting motif located at the C-terminus of the encoded polypeptide and a lysine/arginine rich motif that targets the protein to the plasma membrane (Wang and Chory, 2006; Wang et al., 2014; Novikova et al., 2021). *BKI1* is a single copy gene in Arabidopsis, maize, rice, Brachypodium, and Setaria; however, there are four copies in soybean. The Arabidopsis, rice, and Setaria genes are expressed in most tissues tested with high expression in inflorescences. The maize gene, Zm00001eb315000, and two soybean genes, Glyma.04G038100 and Glyma.06G039100, are expressed high in mature inflorescences as well as young seedling tissues (Figure 9A). This expression pattern difference could indicate a diversification of function in maize and soybean for the *BKI1* genes compared to Arabidopsis, rice, Brachypodium, and Setaria.

**Figure 9.**
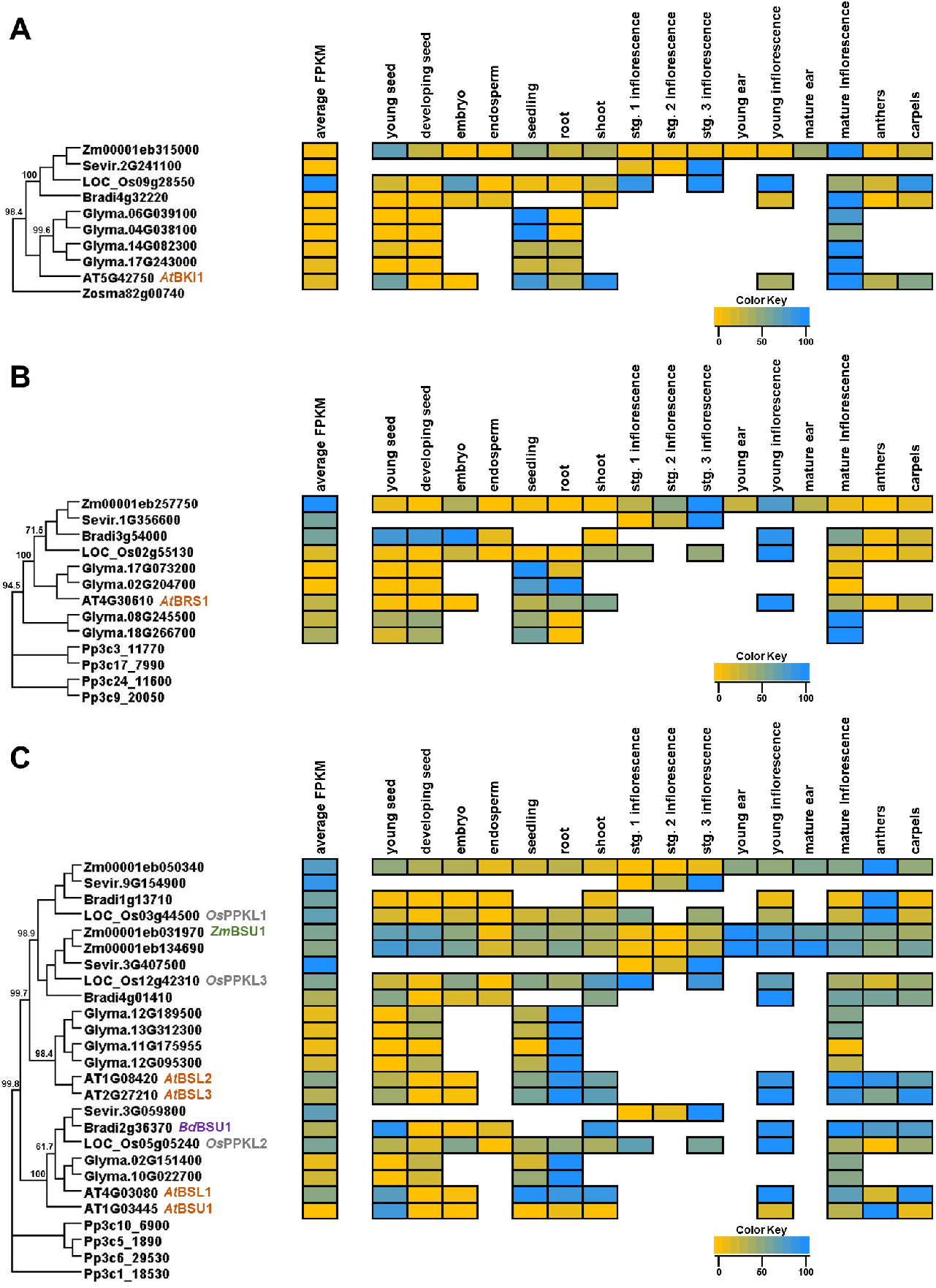
Phylogeny and transcript abundance across different tissues of BRI KINASE INHIBITOR1 (BKI1), BRI1 SUPPRESSOR1 (BRS1), and BRI1 SUPPRESSOR (BSU1) families involved in brassinosteroid signal transduction. Maximum-approximate-likelihood phylogenetic trees of (A) BKI1, (B) BSU, and (C) BRS1 amino acid sequences from maize (Zm), Setaria (Sevir), rice (Os), Arabidopsis (AT), soybean (Glyma), Brachypodium (Bradi), and Physcomitrella (Pp) represented as the outgroup. The relative TPM values from re-analyzed publicly available datasets from different developmental tissues are represented as a heatmap depicted next to each respective gene within the family. The yellow color indicates the lowest individual transcript abundance in each tissue and the dodgerblue color indicates the highest abundance in a given tissue. To compare transcript abundance across genes and species within the family, the average FPKM was determined across all tissues analyzed and is presented in the average FPKM column (left) with the same color distribution as previously described. See Supplemental Tables 1-4 for description of tissue types.

Phenotypic effects of knocking down *At*BKI1 were tested by developing RNAi lines. *AtBKI1*-RNAi lines had longer hypocotyls than wild-type controls when grown in the dark. No other phenotypic effects on plant growth were described. Overexpression of *AtBKI1* in Arabidopsis resulted in severely dwarfed plants resembled *Atbri1* weak mutants. *AtBKI1* overexpression lines were also hyposensitive to BL and hypersensitive to brassinazole (BRZ) treatment and accumulated *AtCPD* and *AtDWF4* transcripts. Increased transcript abundance of *AtCPD* and *AtDWF4* was not, however observed in *AtBKI* overexpressing lines treated with BL (Wang and Chory, 2006), indicating that feedback promotion of BR biosynthetic genes was still dependent on active BR levels in the plant.

In addition to its interaction with BRI1, *At*BKI1 was also shown to interact with 14-3-3 proteins during BR signaling (Wang et al., 2011). 14-3-3 proteins have been shown to regulate BR signaling in both Arabidopsis and rice by inhibiting the nuclear localization of the BES1 and BZR1 transcription factors in the absence of active BRs (Bai et al., 2007; Gampala et al., 2007; Ryu et al., 2007). The C-terminal portion of *At*BKI1 was shown to interact with 14-3-3 proteins to inhibit interaction with *At*BES1/*At*BZR1 and overexpression of 14-3-3 proteins could suppress the inhibitory effect on plant growth in *AtBKI1* overexpression lines (Wang et al., 2011). *AtBKI1* was additionally shown to act in the ERECTA (ER) pathway to regulate plant architecture; it interacts with ER protein family members and inhibits their kinase activity. Lines doubly overexpressing *AtBKI1* and mutant for *Ater* exhibited a more severe dwarf phenotype than either single mutant, as was similarly observed in *Atbri1Ater* double mutants. Furthermore, it was shown that *Atbki1* mutants had a more erect pedicle angle than wild-type controls and could suppress the mutant *Ater* phenotype of less upright, or bent down, pedicels. Thus, *At*BKI1 and *At*ER work together to regulate pedicel orientation in Arabidopsis inflorescences independent of downstream BR signaling (Wang et al., 2017). The *BKI1* genes have not been studied in soybean, maize, rice, Brachypodium, or Setaria.

#### BRI1 SUPPRESSOR1 (BRS1)

The *BRS1* gene encodes a serine carboxypeptidase that functions in BR signaling (Li et al., 2001). BRS1 is secreted from the cell (Zhou and Li, 2005) and its carboxypeptidase activity is necessary for function (Li et al., 2001); however, no interacting partner of BRS1 in BR signaling is known. The *BRS1* gene is single copy in Arabidopsis, maize, rice, Brachypodium and Setaria, and there are four copies in soybean; however, the *BRS1* genes are part of a larger family of serine carboxypeptidases. The highest expression is found in developing inflorescences in all species tested. In Arabidopsis and soybean, expression of *BRS1* is more abundant in other tissues than in maize and rice (Figure 9B). The first mutant *BRS1* mutant described was *Atbrs1-1D* in Arabidopsis, a dominant mutant that suppressed *Atbri1-5* weak mutant phenotypes, including inflorescence length, rosette size, and days to flowering. *Atbrs1-1D* was unable, however, to suppress the flower number and branch number traits affected by *Atbri1-5* (Li et al., 2001). *Atbrs1* loss of function mutants had no visible phenotypes. Overexpressing *BRS1* did increase the number of lateral roots in different accessions and BR signaling mutants (Deng et al., 2017). Further work is necessary to discern the function of *BRS1* in soybean, maize, rice, Brachypodium and Setaria (Liu et al., 2015; Li et al., 2016).

### Brassinosteroid Signal Transduction

#### BRI1 SUPPRESSOR (BSU1)

The *BSU1* gene encodes a phosphatase that acts downstream of BRI1. The BSU1 protein contains a kelch-repeat domain near its N terminus end and a phosphatase domain at the C terminus (Mora-Garcia et al., 2004). Arabidopsis contains three additional homologs of *BSU1* (Figure 9C); *AtBSU1* and *AtBSL1* genes are contained in chromosomal tandem repeats, as are *AtBSL2* and *AtBSL3* (Mora-Garcia et al., 2004). There are six copies of *BSU1*-like genes in soybean. In maize, rice, Brachypodium, and Setaria there are only three copies of *BSU1*-like genes. Interestingly, the subclade with *AtBSU1* and *AtBSL1* includes one rice, Brachypodium, and Setaria gene, but no maize genes. All three maize genes, as well as the other two rice, Brachypodium, and Setaria genes, are in the same subclade as *AtBSL2* and *AtBSL3*. *AtBSU1* is most highly expressed in mature inflorescences, anthers, and young seeds. The *AtBSL1*, *AtBSL2*, and *AtBSL3* transcripts are more evenly distributed in their expression across all tissues analyzed with very low expression in developing seeds and embryos. The six soybean genes are highest expressed in root tissue. Two maize genes, *ZmBSU1* and Zm00001eb134960, are most highly expressed in developing inflorescences and seeds. Another maize gene, Zm00001eb050340, has the highest expression in anthers, like *AtBSU1*. The *OsPPKL1* also has its highest expression in anthers, while *OsPPKL2* and *OsPPKL3* have a more even distribution, and highly similar transcript abundance, across all tissues. All three Setaria genes are highest expressed in stage 3 inflorescences, whereas Sevir.3G059800 has the lowest average expression and is in the same subclade as *At*BSU1 and *At*BSL1 (Figure 9C). Overall, complex phylogenetic and expression pattern relationships suggest substantial functional diversification of *BSU1* family members across species.

The first *AtBSU1* mutant was identified in *Atbri1* mutant suppressor screens, as for the identification of *Atbrs1*. A dominant gain of function allele of the *AtBSU1* gene suppressed *Atbri1*-induced phenotypes affecting leaf and inflorescence development. The study showed that *At*BSU1 was a functional phosphatase that could dephosphorylate *At*BES1 by *At*BIN2-dependent dephosphorylation (Mora-Garcia et al., 2004; Kim et al., 2009). Moreover, *At*BSU1 localized to the cytoplasm could more effectively both suppress *Atbri1-5* mutant phenotypes and dephosphorylated *At*BES1, as compared to nuclear localized *At*BSU1 (Ryu et al., 2010). Loss of function single mutants *Atbsu1*, *Atbsl1*, *Atbsl2*, or *Atbsl3* had no noticeable phenotypes. Expression patterns of the close homologs *AtBSL2* and *AtBSL3* were almost identical, while *AtBSL2* had a higher average expression across tissues (Figure 9C). The double mutants of *Atbsl2* and *Atbsl3*, however, exhibited strong morphological phenotypes. *Atbsl2Atbsl3* double mutant seedlings had epinastic growth with fused cotyledons, but the elongation of dark-grown double mutant seedlings was unaffected. *Atbsl2Atbsl3* double mutants also showed both male and female sterility and produced fasciated inflorescences with less developed vascular bundles. Thus overall, phenotypes observed in *Atbsl2Atbsl3* double mutants did not resemble canonical BR-related mutants, suggesting that *AtBSL1*, *AtBSL2* and *AtBSL3* control plant development independent of *AtBSU1*. Only *AtBSU1* could change the abundance or phosphorylation of canonical BR signaling proteins indicating that this is the only protein within the family that directly acts in the BR signaling pathway (Maselli et al., 2014).

Among other BR signaling pathway components studied in Arabidopsis, the *At*BSU1 protein was also shown to be activated by CONSTITUTIVE DIFFERENTIAL GROWTH1 (*At*CDG1; AT3G26940) in the BR signaling cascade. The *AtCDG1* gene encodes a RLCKVII-subfamily protein kinase localized to the plasma membrane (Muto et al., 2004; Kim et al., 2011). A dominant mutant allele and overexpression of *AtCDG1* resulted in abnormal and epinastic growth and plants were unresponsive to BL treatment effects on hypocotyl growth (Muto et al., 2004). Additionally, the *At*CDG1 and *At*CDG1-like1 (*At*CDL1; AT5G02800) proteins were identified because they were activated by hyperphosphorylated *At*BRI1.

The *At*BR-SIGNALING KINASE1 (*At*BSK1) protein was also shown to be activated by *At*BRI1 and to interact with *At*BSU1. BSK1, a member of the receptor-like cytoplasmic kinase XII family, is localized to the plasma membrane and is phosphorylated by BRI1 upon BR treatment (Tang et al., 2008). There are three homologs in this gene family in Arabidopsis: *BSK1*, *BSK2*, and *BSK3* (Tang et al., 2008). A single knock-out mutant of *Atbsk1* showed no obvious phenotype suggesting redundancy in the pathway in activating *At*BSU1 function, possibly between the BSKs and/or CDG1 (Tang et al., 2008; Kim et al., 2009; Kim et al., 2011). One *BSK* gene in rice, *OsBSK1-1*, was shown to be highly expressed in the leaf joint and overexpression resulted in enhanced BR phenotypes, including increased tillering, flag leaf length, and a less upright leaf angle (Tian et al., 2023). The *Osbsk2* mutant was also identified and showed similar phenotypes to that of *Osbsk1* (Yin et al., 2022). More research is necessary to identify the function of these genes in other species including soybean, Brachypodium, rice, maize and Setaria.

In rice, a strong effect QTL for grain length, qGL3, was identified for which *OsPPKL1* was the underlying, causative locus (Hu et al., 2012; Qi et al., 2012; Zhang et al., 2012). Overexpression of *Os*PPKL1 reduced grain length indicating it negatively regulates growth, and the Kelch domain was necessary for its function to control grain length. By overexpression *Os*PPKL3 also negatively regulated grain length, while *Os*PPKL2 regulated it positively. These contrasts suggest distinct, antagonistic functions of *Os*PPKL2 versus *Os*PPKL1 and *Os*PPKL3 in controlling grain size (Zhang et al., 2012), although expression patterns of these three rice genes are similar across different tissue types (Figure 9C). Recently, CRISPR-*Cas9* was employed to putatively knock out the function of *OsPPKL1*, *OsPPKL2* and *OsPPKL3*. In the single mutants, the only discernable phenotype observed under field or lab conditions was enhanced root elongation in *Osppkl2* when grown on agar medium. This phenotype was more severe in *Osppkl1; Osppkl2* double mutants, suggesting functional overlap with *OsPPKL1* in the root. However, *Osppkl1* did not show altered sensitivity in root growth under excess BR application. Under field conditions, the *Osppkl2; Osppkl3* double mutants had a reduction in plant height, leaf angle, tiller number, and grain size. Double mutants of *Osppkl1; Osppkl3* and the triple mutant were unable to be recovered suggesting that these *OsPPKL1* and *OsPPKL2* may share an essential function(s) (Liu et al., 2021). *Os*PPKL1 was also shown to inhibit the cytokinin phosphorelay cascade and mutation of the specific D364 amino acid resulted in a semi-dominant mutant with larger grain size. This result was independent of the phosphatase domain necessary for BR signaling (Liu et al., 2021). These results exhibit the highly divergent functions of this gene family and the necessity of conducting these comparisons across species. Additional work is needed to identify the function and developmental consequence of *BSU1-*like genes in soybean, maize, Brachypodium, and Setaria.

### Regulation of Brassinosteroid Transcriptional Response

#### BRASSINOSTEROID INSENSTIVE2/GSK3/SHAGGY-LIKE PROTEIN KINASE1 (BIN2/GSK)

The *AtBIN2* gene encodes a GLYCOGEN SYNTHASE KINASE3 (GSK3) -like protein that acts as a negative regulator of BR signaling. BIN2 activity is directly regulated by CDG1 and BSU1 (Kim et al., 2009). The *GSK3* subfamily has ten genes in Arabidopsis that cluster into five separate subclades (Dornelas et al., 1998; Youn and Kim, 2015). Soybean has 22 *GSK3*-like genes, maize has 13, rice has nine, Brachypodium has eight, and Setaria has eight. One of the Arabidopsis and soybean subclades, containing *AtBIN2* along with *AtBIL1* and *AtBIL2*, clusters in the gene tree (Figure 10) as a sister group to five maize genes, four rice genes, three Brachypodium genes, and three Setaria genes. Those grass genes resolve into two additional sister subclades wherein maize, rice and Setaria are represented in all three subclades while Brachypodium is represented in only one (Figure 10). Transcripts of *AtBIN2*, *AtBIL1* and *AtBIL2* are expressed most highly in young inflorescences and carpels and showed high expression in seedling shoots and roots and young seeds. The soybean, maize, rice, Brachypodium, and Setaria genes within this subclade also have their highest transcript expression levels in inflorescences, except for two soybean genes (Glyma.12G212000 and Glyma.13G228100) that have their highest expression in developing seeds (Figure 10).

**Figure 10.**
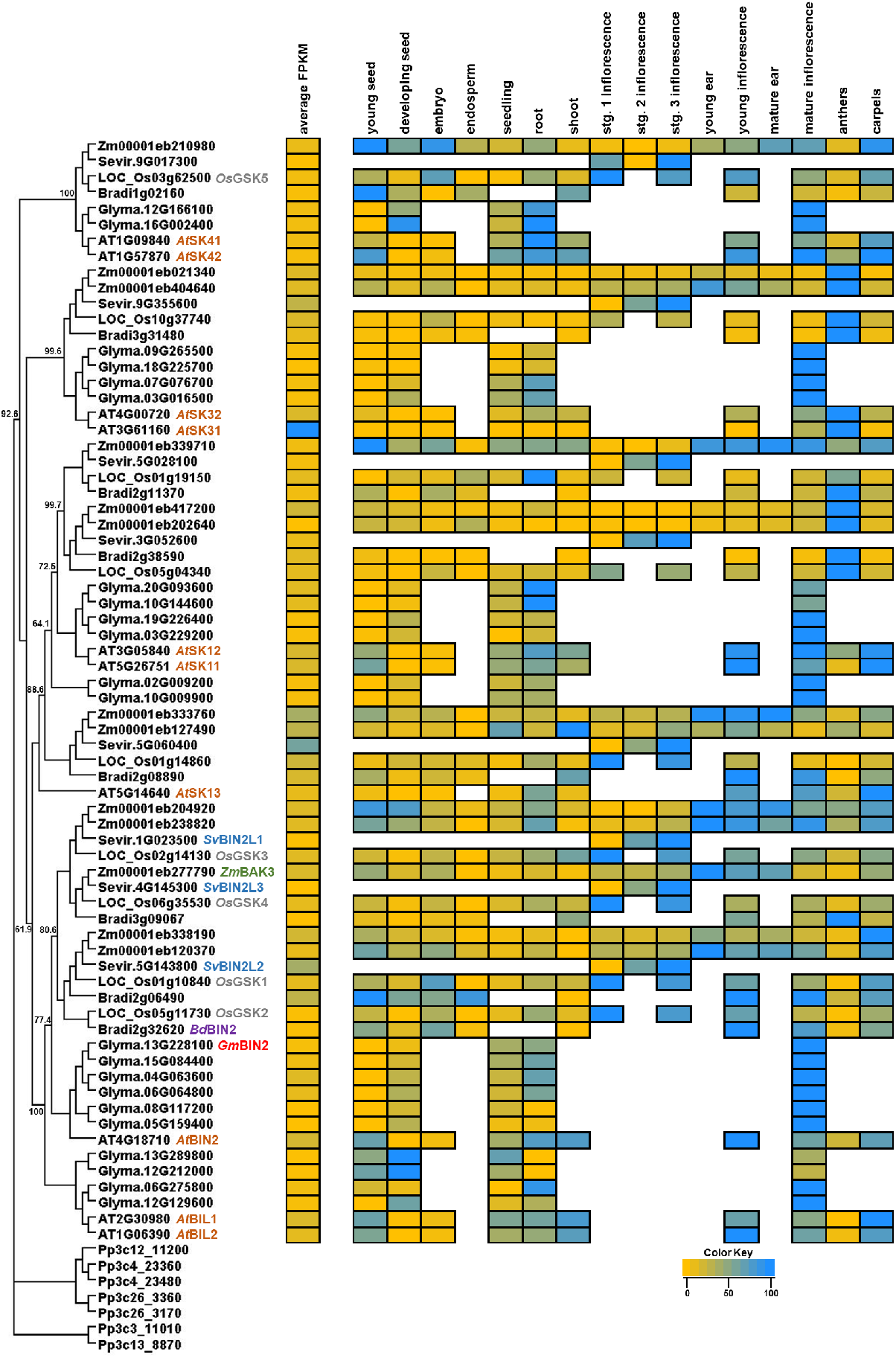
Phylogeny and transcript abundance across different tissues of the BRASSINOSTEROID INSENSTIVE2/GSK3/SHAGGY-LIKE PROTEIN KINASE1 (BIN2/GSK) family involved in brassinosteroid signal transduction. Maximum-approximate-likelihood phylogenetic tree of BIN2/GSK amino acid sequences from maize (Zm), Setaria (Sevir), rice (Os), Arabidopsis (AT), soybean (Glyma), Brachypodium (Bradi), and Physcomitrella (Pp) represented as the outgroup. The relative TPM values from re-analyzed publicly available datasets from different developmental tissues are represented as a heatmap depicted next to each respective gene within the family. The yellow color indicates the lowest individual transcript abundance in each tissue and the dodgerblue color indicates the highest abundance in a given tissue. To compare transcript abundance across genes and species within the family, the average FPKM was determined across all tissues analyzed and is presented in the average FPKM column (left) with the same color distribution as previously described. See Supplemental Tables 1-4 for description of tissue types.

The *AtSK13* subclade is similarly complex with inclusion of two additional Arabidopsis genes *AtSK11* and *AtSK12* and several soybean, Setaria, Brachypodium, maize, and rice genes. Increased copy number in maize and soybean reflects a likely duplication of this gene specific to both species. *AtSK11* and *AtSK12* transcripts accumulate across tissues in a similar manner to the *AtBIN2*/*BIL* transcripts with their highest expression in inflorescences and carpels. These genes also have high transcript abundance in seedling tissues and also accumulate in young seeds (Figure 10).

Transcripts of *AtSK31* and *AtSK32* are most highly expressed in anthers with the *AtSK31* transcripts having the highest average FPKM across all tissues. Related paralogs in other species also have their highest expression in inflorescences (Figure 10). Arabidopsis *AtSK41* and *AtSK42* genes form a final subclade that includes a single gene each from maize, rice, Brachypodium and Setaria; but there are two soybean genes like in Arabidopsis, showing a diversification between eudicots and monocots. *AtSK41* and *AtSK42* have relatively uniform transcript abundance across all tissues with their lowest abundance in developing seeds and embryos. Related maize, rice, Brachypodium, and Setaria genes show similar transcript abundance across tissues compared to the Arabidopsis genes, except the grass genes have relatively higher expression in developing embryos (Figure 10).

*BIN2* was first characterized in Arabidopsis by identification of the semi-dominant mutant *Atbin2* that resembled BR deficient mutants and was hyposensitive to BL treatment for root seedling elongation phenotypes (Li et al., 2001). The determination that *BIN2* negatively regulates BR responses was further supported by observations that overexpression of *Bd*BIN2 in Arabidopsis which showed a dwarf phenotype (Corvalán and Choe, 2017).

*Atbin2* mutants were also hypersensitive to ABA treatment whereas their roots were significantly shorter than wild-type plants, as seen for *Atbri1* and BR-deficient mutants (Clouse et al., 1996; Ephritikhine et al., 1999), therefore suggesting crosstalk between BR and ABA to regulate root growth in Arabidopsis. In investigations of BR and auxin crosstalk, root growth of *Atbin2* mutants was mildly hyposensitive to synthetic auxin treatment (Choe et al., 2002), but not to natural auxin treatment (Li et al., 2001). The infertility phenotype of dominant *Atbin2* mutants could be partially rescued with exogenous application of the synthetic auxin naphthylacetic acid (NAA) (Li et al., 2020), suggesting a complex interaction between auxin and BRs to regulate plant development. A key function of the BIN2 protein kinase is to phosphorylate BES1/BZR1 transcription factors to keep them from entering the nucleus (He et al., 2002). Dominant *Atbin2* alleles contain mutations in the TREE motif, part of the BIN2 catalytic domain, resulting in higher kinase activity (Choe et al., 2002; Li and Nam, 2002; Yan et al., 2009). *Atbin2* mutants were unable to regulate feedback of BRs as the semi-dominant mutants had higher levels of endogenous BRs, especially downstream substrates (Choe et al., 2002).

The *At*BIN2 protein has also been shown to regulate the function of other proteins and pathways in BR signaling, in addition to those downstream of BES1/BZR1 transcription factors. For example, BIN2 regulates cell elongation independent of BES1/BZR1 transcription factor-coupled pathways by phosphorylating AUXIN RESPONSE FACTOR2 (ARF2), PHYTOCHROME INTERACTING FACTOR4 (PIF4), and CESTA transcription factors (Vert and Chory, 2006; Bernardo-Garcia et al., 2014; Khan et al., 2014). Regulation of root development by *At*BIN2 has been shown by its ability to phosphorylate and regulate the function of *At*ARF7, *At*TRANSPARENT TESTA GLABRA1 (*At*TTG1), and *At*ENHANCER OF GLABRA3 (*At*EGL3) (Cheng et al., 2014; Cho et al., 2014). *At*BIN2 protein can also negatively regulate cellulose synthesis by directly phosphorylating *At*CELLULOSE SYNTHASE A1 and inhibiting its function, and the dominant *Atbin2* mutants have lower cellulose content (Sanchez-Rodriguez et al., 2017). *At*BIN2 is also involved in drought response, where phosphorylation of *At*RD26 results in increased activation and transcriptional regulation of drought-responsive genes (Jiang et al., 2019). *At*BIN2 has an additional role in directly regulating the balance between salt stress and growth recovery in Arabidopsis. The *At*BIN2 protein was able to phosphorylate and inhibit function of *At*SALT OVERLY SENSITIVE2, thus inhibiting the salt dependent stress response (Li et al., 2020). These findings were supported by overexpression of *Gm*BIN2 in Arabidopsis and soybean hairy roots that showed a greater drought and salt tolerance (Wang et al., 2018). Taken together, the activity of BR signaling through BIN2 can regulate multiple plant growth and stress responses independent of the canonical BR response pathway transcription factors.

To date, *AtBIN2* loss of function mutants have not been recovered from BR forward genetic screens. To test the possible redundancy of *AtBIN2* with other GSK3 kinase genes in Arabidopsis, *Atbin2* loss of function alleles were first developed by mutagenizing plants containing a weak dominant *Atbin2* allele and screening for intragenic suppressors that eliminated the dominant phenotype. Several loss of function *Atbin2* alleles showed no obvious phenotypes as compared to wild-type siblings (Yan et al., 2009), likely due to redundancy of GSK3 proteins. However, the loss of function allele *Atbin2-3* did partially recover the *Atbri1-5* dwarf phenotype and single mutants of *Atbin2-3* were slightly hyposensitive to BL treatment for root and hypocotyl growth. T-DNA mutant lines of both *Atbil1* and *Atbil2* were characterized to test the hypothesis of possible redundancy among GSK3 proteins. The single and double mutants between *Atbil1* and *Atbil2* exhibited no visible phenotypes under normal growth conditions, but like *Atbin2-3* were less sensitive to BR and BRZ application, suggesting redundancy between these three genes. The single *Atbin2*, *Atbil1*, and *Atbil2* loss of function mutants were also hyposensitive to ABA-induced root inhibition due to their interaction with *At*SNF1-RELATED KINASE2S (Cai et al., 2014). Beyond its role in BR signaling, *At*BIL1 also functions in auxin and cytokinin signaling through phosphorylation of *At*MONOPTEROS to transcriptionally activate negative regulators of cytokinin signaling and control vascular cambium proliferation (Han et al., 2018). The double and triple mutants of *Atbil1* and/or *Atbil2* with *Atbri1-5* could not recover the *Atbri1*-5 dwarf phenotype. However, when making mutant combinations with *Atbil1* and/or *Atbil2* with *Atbin2-3* and *Atbri1*, the suppression of the *Atbri1-5* phenotype was greatly enhanced indicating further redundancy of these loci in Arabidopsis. Interestingly, triple mutants for *Atbin2-3*, *Atbil1*, and *Atbil2* were still responsive to BRs, suggesting other GSK3s could also be functioning in BR signaling (Yan et al., 2009).

Consistent with this hypothesis, overexpressed *At*SK32 protein was shown to directly phosphorylate *At*BES1/*At*BZR1 transcription factors and regulate their transport into the nucleus (Kim et al., 2009; Rozhon et al., 2010). Furthermore, recessive mutations of *Atsk32* have reduced hypocotyl elongation and altered floral organ cell size by influencing the transcriptional levels of *AtXYLOGLUCAN ENDOTRANSGLYCOSYLASES* (*AtXET*s) (Claisse et al., 2007). The *At*SK31 protein is a component of the CONSITITUTIVE PHOTOMORPHOGENIC9 (COP9) complex that represses photomorphogenesis and induces skotomorphogenesis (Wei et al., 1994; Chamovitz et al., 1996; Staub et al., 1996). Mutants of the COP9 complex, including *Atsk31* (Wei et al., 1994), have phenotypes that resemble BR deficient mutants when grown in the dark (Karniol and Chamovitz, 2000). The COP9 signalosome regulates many developmental pathways in Arabidopsis (Karniol and Chamovitz, 2000), but no direct connection between the COP9 complex and BRs has been made.

The *At*SK11 and *At*SK12 proteins have also been shown to be involved in BR signaling by physically interacting with BES1/BZR1 transcription factors via yeast-2-hybrid and Co-IP assays (Kim et al., 2009; Tang et al., 2011). Both *At*SK11 and *At*SK12 transcripts are expressed in developing flowers, based on *in situ* hybridization, with their expression highly overlapping in inflorescence and floral meristems. Transgenic antisense constructs of *At*SK11 and *At*SK12 lead to abnormal flower development with increased sepal, petal, and flower bud numbers and increased the number of floral meristems as compared to wild-type siblings. Antisense plants also have altered gynoecium apical-basal patterning and a larger gynophore at the basal end of the gynoecium. The antisense transgenes did not, however, affect flowering time or the pattern of cell division. (Dornelas et al., 2000). By contrast, the *At*SK12 protein recently was shown to regulate flowering by degrading CONSTANS via direct phosphorylation. Recessive mutants of *Atsk12* flowered earlier and had fewer rosette leaves (Chen et al., 2020), suggesting the antisense lines were knockdowns with residual *At*SK function. Further studies are needed on recessive mutants of *AtSK11* and *AtSK12* to identify the direct effects these loci have on the BR signaling pathway.

Among additional regulators of *At*BIN2, the F-Box E3 ubiquitin ligase *At*KINK SUPPRESSED IN BZR1-1D (*At*KIB1) was also shown to regulate *At*BIN2 function via ubiquitination and subsequent targeting for proteasomal degradation. *At*KIB1 could also inhibit *At*BIN2 by interfering with substrate access of other interacting proteins including the *At*BES1/*At*BZR1 transcription factors. The *At*KIB1 protein could physically interact with *At*BIN2 as shown by *in vivo* Co-IP experiments and the C-terminal end of *At*KIB1 was sufficient to interact with *At*BIN2 via yeast two-hybrid assays (Zhu et al., 2017). Furthermore, the C-terminal end of *At*KIB1 was also shown to physically interact with *At*BIL1, *At*BIL2, *At*SK11, *At*SK12, and *At*SK13 in yeast two-hybrid assays (Zhu et al., 2017), further implicating these proteins in BR signaling and suggesting a high overlap of redundancy among these members of the GSK3 family. In rice, it has also been shown that the U-box ubiquitin ligase *Os*TUD1 is able to physically interact with *Os*GSK2 and ubiquinate the protein and inhibit its function (Liu et al., 2023).

The *AtS*K41 and *At*SK42 proteins show high amino acid sequence similarity to *At*BIN2 (Jonak and Hirt, 2002), however their functions in plant development and/or BR signaling have not yet been characterized. Transcript abundance of *At*SK41 and *At*SK42 are relatively lower than the other eight GSK3 genes across most tissues (Figure 10) (Charrier et al., 2002). The *At*SK42 transcriptional levels increased after sodium chloride and polyethylene glycol treatments, whereas *At*SK41 was unresponsive (Charrier et al., 2002). Further studies are needed to identify specific functions of *At*SK41 and *At*SK42.

The maize BIN2/GSK3 genes have been investigated through the development of RNAi lines (Kir, 2015). These lines redundantly targeted all *Zm*BIN2/*Zm*GSK3-like transcripts by using the full-length cDNA of Zm00001eb204920. This resulted in lower transcriptional accumulation of all loci tested in leaf tissue; however, Zm00001eb338190 and Zm00001eb417200 were not expressed in this tissue (Kir, 2015) and Zm00001eb20224460, Zm00001eb404640, and Zm00001eb021340 were not tested (Figure 10). Therefore, RNAi phenotypes could not be assigned to a particular locus. RNAi lines showed variable severity of phenotypes ranging from moderate to severe. All events showed a reduction in plant height due to a relatively uniform reduced internode length throughout development (Kir, 2015). These knockdown phenotypes are surprising given that combinations of *BIN2*/*GSK3* mutant alleles in Arabidopsis did not show visible phenotypes (Yan et al., 2009) and underline the necessity of developing higher order mutant combinations in Arabidopsis to alleviate phenotypic masking due to functional redundancy of these homologs. The *Zmbin2*-RNAi lines flowered later than wild-type siblings and inflorescences were elongated and had a lower spikelet density with many barren tips, suggesting a reduction in spikelet pair initiation and/or development. Interestingly, *Zmbin2*-RNAi lines also showed maternally increased kernel number and embryo size (Kir, 2015). Similar to Arabidopsis mutants, *Zmbin2*-RNAi lines had reduced root growth (Yan et al., 2009). However, the *Zmbin2*-RNAi lines were hyposensitive to BL treatment (Kir, 2015), dissimilar to Arabidopsis where *Atbin2-3* mutants were hypersensitive to BL effects on root growth (Yan et al., 2009). To identify the individual effects each BIN2/GSK3 locus has in maize, individual mutants and then different mutant combinations will need to be developed.

The rice *Os*GSK1, *Os*GSK2, *Os*GSK3, and *Os*GSK4 genes have been well described. Development of overexpression, RNAi and CRISPR/*Cas9* lines has enabled comparison of differential contributions of these loci to control plant development. Overexpression of *Os*GSK2 had no obvious effect on plants, but overexpression of mutated forms with presumed hyperactive kinase activity resulted in strong BR-deficiency phenotypes including reduced plant height, tillering and growth of leaf sheathes and blades, as well as upright leaf angle, delayed flowering time, erect panicles, some sterility, and reduced grain size (Tong et al., 2012). RNAi-lines targeting *Os*GSK2 resulted in reduced transcription levels of *Os*GSK1, *Os*GSK2, *Os*GSK3, and *Os*GSK4. These RNAi-lines did not affect plant height but did exhibit less upright leaves that were longer and narrower than wild-type siblings. The grain length was also increased in these transgenic lines (Tong et al., 2012; Liu et al., 2021). To investigate possible redundancy of the BIN2/GSK3 loci in rice, independent CRISPR/*Cas9* lines were developed for all four genes. *Osgsk2* single mutants and double mutants with *Osgsk1*, *Osgsk3*, or *Osgsk4* were significantly taller and had more upright leaves than wild-type siblings. No other single mutant phenotypes were reported. Triple mutants between *Osgsk1*, *Osgsk2*, and *Osgsk3* had no significant effect on plant height, but had a less upright leaf angle. Triple mutant combinations between *Osgsk2*, *Osgsk3*, and *Osgsk4*, as well as quadruple mutants of all four loci, reduced plant height made leaves less upright as compared to wild-type siblings. All mutant combinations tested, as well as *Osgsk2* single mutants, increased grain size primarily by increasing grain length, like *Osgsk* RNAi-lines. The phenotypes described for the *Osgsk* CRISPR/Cas9 and *Osgsk* RNAi-lines closely resembled those of the maize *Zmbin2*-RNAi lines suggesting a conservation of function between maize and rice.

The *Os*GSK2 protein has also been shown to directly phosphorylate another positive regulator of BR signaling, the GRAS transcription factor encoded by *OsDWARF AND LOW-TILLERING* (*OsDLT*) (Tong et al., 2012). Loss-of-function *Osdlt* mutants have a reduced stature, upright leaves, and reduction of tillering. These mutants are also hyposensitive to BL treatment in coleoptile length and lamina joint angle assays (Tong et al., 2009). Overexpression of *OsDLT* could suppress the phenotypes of *OsGSK2* overexpression lines indicating that *OsDLT* functions downstream *OsGSK2* in rice (Liu et al., 2021). Another GRAS transcription factor in rice, WIDE GRAIN3 (WG3), was shown to physically interact with *Os*DLT to promote BR signaling and increase grain size (Chen et al., 2022). *Os*DLT has also been implicated in GA metabolism as loss of function *Osdlt* mutants have reduced GA levels, reduced α-amylase activity in seeds, and upregulated GA biosynthetic gene transcripts (Li et al., 2010). It is yet unclear whether the effects on GA-metabolism in *Osdlt* mutants are independent of BR signaling or result from crosstalk between these two hormones.

Further work is needed to investigate the function of, and developmental consequences of altering, other *BIN2*/*GSK3* loci in rice (Figure 10). The function of *BIN2*/*GSK3* loci in Setaria and Brachypodium have not been greatly investigated to date. The most closely related loci to *AtBIN2*, i.e. *SvBIN2L1*, *SvBIN2L2* and *SvBIN2L3* (Figure 10), had only slightly reduced transcript levels compared to wild-type siblings in the biosynthetic mutant *Svbsl1* and did not meet the genome-wide corrected *p*-value of 0.05 (Yang et al., 2018). More work is needed to develop genetic resources to identify the function of these genes in Setaria and Brachypodium.

#### BRI1-EMS-SUPPRESSOR1/BRASSINAZOLE-RESISTANT1 (BES1/BZR1)

The *BES1/BZR1* genes encode beta-helix-loop-helix (bHLH) type transcription factors that positively regulate BR signaling (He et al., 2002; Wang et al., 2002; Zhao et al., 2002). There are six *BES1*/*BZR1* like genes in Arabidopsis, ten in soybean, seven in maize, four in rice, six in Brachypodium, and three in Setaria (Figure 11). *AtBES1*, *AtBZR1*, *AtBEH1*, and *AtBEH2* are in the same subclade along with a single gene from rice and Setaria, two in maize, three in Brachypodium and six in soybean. *AtBEH3* and *AtBEH4* belong to a second subclade that contains four soybean genes, two genes from maize, Brachypodium and rice, and two genes from Setaria. Three maize genes comprise another, separate subclade. *AtBES1* and *AtBZR1* show similar expression profiles across tissues with *AtBES1* being most highly expressed in seedling shoots and *AtBZR1* most highly expressed in carpels. *AtBES1* has the highest average FPKM value across all tissues in Arabidopsis. *AtBEH1* is also most highly expressed in seedling shoots, but interestingly it has low expression in mature inflorescences, anthers, and carpels. *AtBEH2* is also highly expressed in young and mature inflorescences (Figure 11). These data suggest variable gene regulation among the closely related Arabidopsis genes in reproductive structures. *AtBEH3* and *AtBEH4* also have similar expression distributions across tissues with their highest expression in young inflorescences. All soybean genes have similar expression to their closest homologs in Arabidopsis (Figure 11).

**Figure 11.**
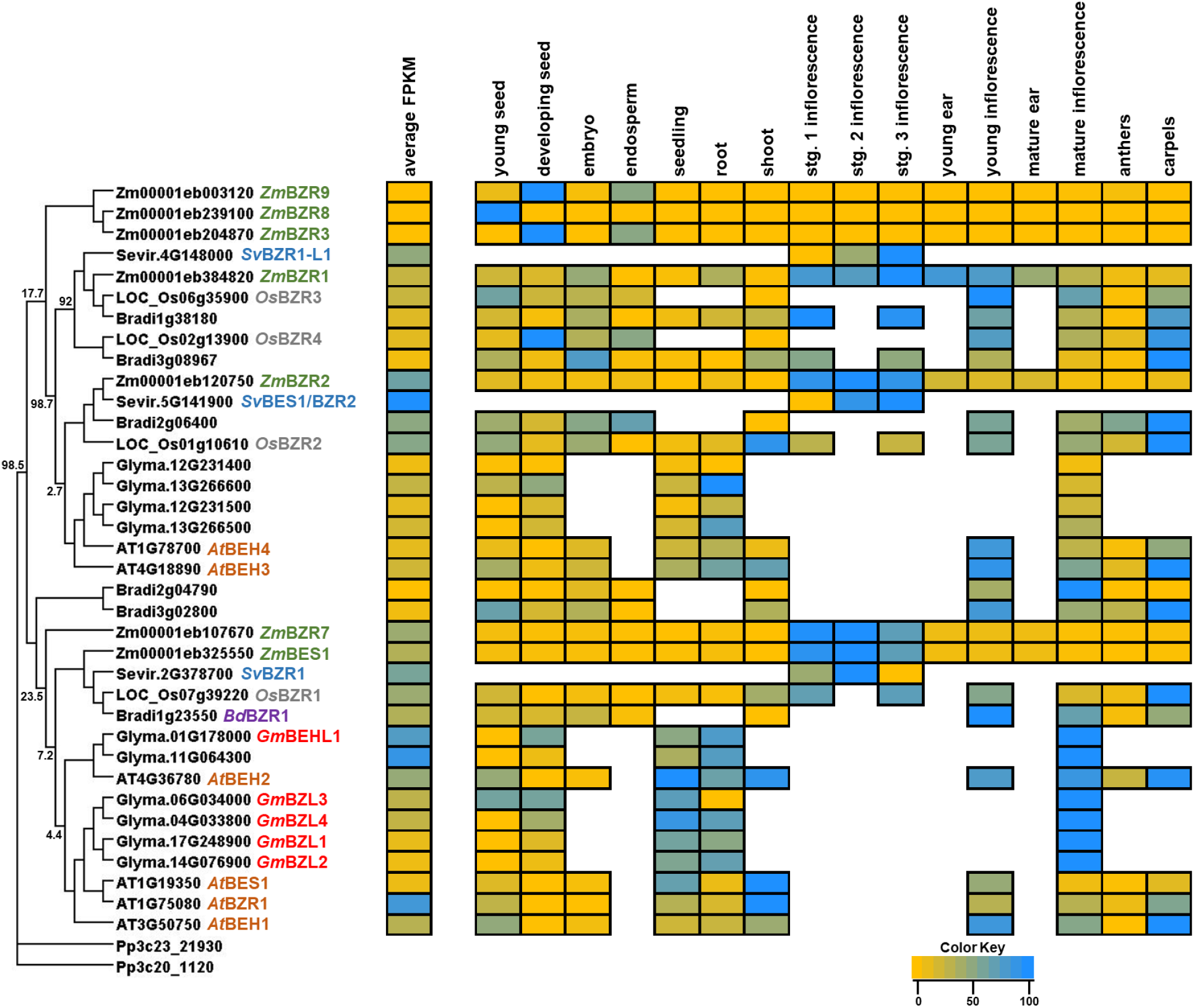
Phylogeny and transcript abundance across different tissues of the BRI1-EMS-SUPPRESSOR1/BRASSINAZOLE-RESISTANT1 (BES1/BZR1) brassinosteroid transcription factor family. Maximum-approximate-likelihood phylogenetic tree of BES1/BZR1 amino acid sequences from maize (Zm), Setaria (Sevir), rice (Os), Arabidopsis (AT), soybean (Glyma), Brachypodium (Bradi), and Physcomitrella (Pp) represented as the outgroup. The relative TPM values from re-analyzed publicly available datasets from different developmental tissues are represented as a heatmap depicted next to each respective gene within the family. The yellow color indicates the lowest individual transcript abundance in each tissue and the dodgerblue color indicates the highest abundance in a given tissue. To compare transcript abundance across genes and species within the family, the average FPKM was determined across all tissues analyzed and is presented in the average FPKM column (left) with the same color distribution as previously described. See Supplemental Tables 1-4 for description of tissue types.

In maize, *ZmBES1* and *Zm*BZR7 are most highly expressed in small, developing tassels. *ZmBZR2* has a higher average FPKM value than *ZmBES1* and *ZmBZR7* which is surprising given its placement in the subclade with *AtBEH3* and *AtBEH4*. *ZmBZR1* and *ZmBZR7* are also highly expressed in developing maize tassels, but *ZmBZR1* is also more highly expressed in young seeds, embryos, and roots relative to the other maize genes expression profiles within subclades with characterized Arabidopsis loci. The three maize genes that comprise their own subclade are most highly expressed in seeds (*ZmBZR8*) and endosperm (*ZmBZR3* and *ZmBZR9*) suggesting a diversification of the role of these three *BZR* genes within maize as well as compared to the other species (Figure 11). However, the very low transcript abundance across all tissues for each of the three genes could indicate they are pseudogenes (Figure 11). The four rice *BZR* transcripts show highest expression in developing inflorescences and carpels, suggesting conservation and possible redundancy of these genes in rice. The three rice transcripts in the subclade with *AtBEH3* and *AtBEH4* are also relatively well expressed in developing embryos compared to other tissues. Brachypodium genes have similar expression to their closest homologs in rice and maize (Figure 11).

The first characterized *BES1*/*BZR1* mutant in Arabidopsis, *Atbrassinazole-resistant1-1D* (*Atbzr1-1D*), was identified as a dominant allele in a screen for mutants insensitive to BRZ, a BR biosynthetic inhibitor that phenocopies BR deficient mutants (Asami et al., 2000; He et al., 2002; Wang et al., 2002; Zhao et al., 2002). *Atbzr1-1D* mutants were also hyposensitive to BL treatment when grown in the dark but had a similar hypocotyl length as compared to wild type when grown on normal media. When grown in the light, the *Atbzr1-1D* mutants were shorter, had more and wider dark green leaves, delayed flowering, and were epistatic to BR biosynthetic and other BR signaling mutants. The *Atbzr1-1D* mutants had reduced levels of downstream BR intermediates as well as a reduction of *At*CPD transcript accumulation indicating a feedback inhibition on BR biosynthesis (Wang et al., 2002). The *At*BZR1, *At*BES1, *At*BEH1, *At*BEH2, *At*BEH3, and *At*BEH4 proteins all contain a GSK3 phosphorylation site and can be directly phosphorylated by *At*BIN2 (Wang et al., 2002; Yin et al., 2002). The *Gm*BEHL1 protein was also shown to interact with *Gm*BIN2 by yeast-2-hybrid and biomolecular fluorescence complementation assays (Yan et al., 2018). Phosphorylation of BES1/BZR1 proteins results in them being localized strictly to the cytosol (He et al., 2002; Wang et al., 2002; Yin et al., 2002; Zhao et al., 2002). Inhibition or shuttling of phosphorylated BES1/BZR1 proteins out of the nucleus requires their association with 14-3-3 binding proteins (Bai et al., 2007; Gampala et al., 2007; Ryu et al., 2007). Mutation of the *At*BIN2 binding site in *At*BZR1 abolished 14-3-3 association and increased *At*BZR1 localization in the nucleus (Gampala et al., 2007). There are multiple copies of 14-3-3 proteins in Arabidopsis and neither knock-out mutations nor RNAi lines resulted in any visible phenotypes, likely due to a redundancy in function across the multiple copies (Gampala et al., 2007). Studies have also shown that the *At*BES1/*At*BZR1 transcription factors can be activated independently of the BR signaling pathway (Zheng et al., 2019; Albertos et al., 2022; Bai et al., 2022; Chen et al., 2022; Shi et al., 2022; Zheng et al., 2022).

The *At*BRZ SENSITIVE SHORT HYPOCOTYL1/*At*BLADE ON PETIOLE1 (*At*BSS1/*At*BOP1) protein was shown to inhibit the nuclear location of *At*BZR1 protein (Shimada et al., 2015). Overexpression of *At*BSS1/*At*BOP1 resulted in reduced nuclear localization of *At*BZR1, while deficient mutants of *Atbss1*/*Atbop1* had increased nuclear localization of *At*BZR1. The loss-of-function *At*BSS1/*At*BOP1 mutants had reduced petioles with ectopic outgrowths, reduced number of rosette leaves, and longer flowering time (Ha et al., 2003). Dominant and overexpression lines of *At*BSS1/*At*BOP1 resulted in strong BR deficient mutant phenotypes (Shimada et al., 2015), further supporting its involvement in inhibiting BR signaling.

Once BRs are present, BIN2 inhibition of BES1/BZR1 transcription factors is removed by subsequent targeting of BIN2 to the 26s proteasome for degradation. Removal of *At*BIN2 results in a dephosphorylation of *At*BES1/*At*BZR1 transcription factors by *At*PROTEIN PHOSPHATASE2A (*At*PP2A) (Di Rubbo et al., 2011; Tang et al., 2011). The unphosphorylated BES1/BZR1 transcription factors can the enter the nucleus to bind and regulate BR responsive genes. The *At*BES1/*At*BZR1 transcription factors were shown to recruit another bHLH transcription factor, *At*BES1 INTERACTING MYC-LIKE1 (*At*BIM1), and bind to E-box motifs to promote transcription of BR responsive genes (Yin et al., 2005). They have also been shown to bind to BR-Response Elements (BRRE) in the promoters of BR responsive genes to inhibit transcription (He et al., 2005). The *At*BES1/*At*BZR1 transcription factors contain an ERF-ASSOCIATED AMPHIPHILIC REPRESSION (EAR) motif near the C-terminal end of the protein that is required for *At*BES1/*At*BZR1 regulation of gene expression (Wang et al., 2013). Deletion of the EAR motif eliminates their ability to repress gene expression and regulate growth and development, but that activity could be recovered by fusing the Groucho/TUP1-LIKE transcriptional corepressor *At*TOPLESS (*At*TPL) to the EAR motif-less *At*BZR1 (Oh et al., 2014). This result indicated that recruitment of *At*TPL by *At*BZR1 was necessary to repress BR regulated gene expression. Quadruple mutants of *Attpl*; *Attopless-related1* (*Attpr1*); *Attpr4*; *Atbzr1-1D* suppressed the *Atbzr1-1D* mutant phenotypes further indicating the necessity of *At*TPL for *At*BZR1 to function properly (Oh et al., 2014). *Attpl* single mutants were temperature sensitive and showed altered apical embryonic phenotypes with a transition of the shoot apical meristem into a root meristem at certain temperatures (Long et al., 2002; Long et al., 2006). Ectopic expression of *At*TPL resulted in less upright leaves and altered inflorescence development in a *At*BZR1/*At*BES1 dependent manner (Espinosa-Ruiz et al., 2017). *At*TPL has also been shown to be involved in jasmonic acid and auxin signaling (Szemenyei et al., 2008; Pauwels et al., 2010), indicating a potential crosstalk between BR signaling and other hormone signaling pathways.

Recessive loss-of-function single mutants *Atbzr1*, *Atbes1*, *Atbeh3* and *Atbeh4* grown in the dark had significantly shorter hypocotyls compared to wild-type siblings; however, the reduction was not as severe as seen in biosynthetic mutants (Lachowiec et al., 2018). In soybean, *Gm*BEHL1 was shown to negatively regulate nodule number, but result in larger nodules (Yan et al., 2018). The *Atbeh1* and *Atbeh2* single mutants did not have significantly different hypocotyl lengths compared to wild type. The *Atbeh4-1* mutant was the only mutant to have a decrease in developmental robustness of hypocotyl growth when grown in the dark and was dependent on *At*BES1 function (Lachowiec et al., 2018). None of the *Atbes1*/*Atbzr1* mutants influenced developmental robustness when grown in the light. There were no other obvious vegetative defects observed in the single mutants nor double, triple, quadruple, or pentuple mutants of *Atbes1*/*Atbzr1* combinations studied, except for the *Atbzr1*;*Atbes1*;*Atbeh1*;*Atbeh3*;*Atbeh4* quintuple mutants that exhibited curled leaves and a semi-dwarf stature (Lachowiec et al., 2018; Chen et al., 2019). Hextuple mutants of all six *Atbes1*/*Atbzr1* loci resulted in severe dwarf phenotypes and resembled severe *Atbri1* recessive mutants. This result indicated a high level of redundancy between all *At*BES1/*At*BZR1 transcription factors to regulate the expression of BR responsive genes to regulate growth and development. Redundancy appears conserved between Arabidopsis and soybean based on mutant analysis of the *GmBZL2* and *GmBZL3* genes (Zhang et al., 2016; Song et al., 2019). Notably, *At*BEH2 is likely the weakest of the six *At*BES1/*At*BZR1 transcription factors, as the pentuple mutant of the other five genes did exhibit a weak phenotype (Chen et al., 2019), even though *At*BEH2 can be phosphorylated and regulated by *At*BIN2 (Rozhon et al., 2010).

Some mutants of *Zm*BES1/*Zm*BZR1 transcription factor genes have been described in maize. Uniform*Mu* (Settles et al., 2007) and CRISPR/Cas9 knock-out mutants were developed for *Zmbes1* and *Zmbzr1* (Wang et al., 2022). Both single mutants were significantly shorter and had more erect leaves than wild-type siblings. There was no effect on leaf length or width for either single mutant. The double mutants between *Zmbes1* and *Zmbzr1* were not more severe than either single mutant for plant height but were additive for leaf angle, resulting in more erect leaves than either single mutant (Wang et al., 2022). This result is surprising as single mutants of BES1/BZR1 in Arabidopsis do not have a light-grown phenotype and suggests a diversification of function in maize. No other canonical BES1/BZR1 mutants have been described in maize. Development of higher order mutant combinations will most likely be necessary to observe a severe phenotype similar to those of biosynthetic mutants (Hartwig et al., 2011; Makarevitch et al., 2012; Best et al., 2016).

A recessive mutant of the *OsDLT* ortholog, *ZmSCARECROW-LIKE28* (*ZmSCL28*), was also developed and the *Zmscl28* mutant was shorter and had more erect leaves than either *Zmbes1* or *Zmbzr1*. Double mutants between *Zmscl28* and *Zmbes1* or *Zmbzr1* were not phenotypically altered from *Zmscl28*, indicating genetic epistasis between *ZmSCL28* and *ZmBES1*/*ZmBZR1*. Yeast one-hybrid experiments also supported this result as *Zm*BES1 and *Zm*BZR1 were able to bind the promoter of *Zm*SCL28 (Wang et al., 2022). Thus, interaction between DLT and BES1/BZR1 transcription factors appears to be conserved between rice and maize.

The *ZmBZR2* gene (Yu et al., 2018) was overexpressed in Arabidopsis where it resulted in larger leaf size and seed size (Zhang et al., 2020). Enlargement of these organs was both proximal-distal and medial-lateral due to larger cell size in the overexpression lines. This locus (v3-GRMZM5G852801/GRMZM6G287292; v4-Zm00001d039439) was not included in our phylogenetic and expression analysis (Figure 11) because it was not annotated in version 5 of the maize genome. The *ZmBZR6* gene has also been molecularly characterized in maize (Yu et al., 2018). The *Zm*BZR6 protein contains a bHLH domain and was predicted to bind to BRRE and E-box motifs, but it also contains a β-amylase (BAM) domain (Sun et al., 2020). The closest Arabidopsis homologs (AT2G45880 (*At*BAM7) and AT5G4530 (*At*BAM8)) also contain a BAM domain in addition to the bHLH domain (Reinhold et al., 2011). Single mutants of *Atbam7* and *Atbam8* showed no discernable phenotypes, but the *Atbam7*; *Atbam8* double mutants did result in slower growth and smaller leaves compared to wild-type siblings. Overexpression of these two loci resulted in larger leaves (Reinhold et al., 2011). Mutants of *ZmBZR6* resulted in smaller seed size, while overexpression of *Zm*BZR6 in rice resulted in larger seed size (Sun et al., 2021). Overexpression in Arabidopsis caused a decrease in ABA sensitivity but enhanced salt and drought tolerance (Sun et al., 2020). These BAM transcription factors were proposed to be involved in an alternative signaling pathway other than BR signaling, even though they may compete for the same genetic targets (Reinhold et al., 2011). Direct control of BAM function by other characterized BR signaling components has not been reported.

The four rice *OsBES1*/*BZR1* genes were characterized using CRISPR/*Cas9*. Among single mutants only *Osbzr2* exhibited a phenotype, including shorter plants, less upright leaves, and suppressed tillering (Liu et al., 2021). Higher order mutant combinations were obtained by combining weak mutant alleles at one locus with stronger alleles at the others. No double or triple, or quadruple mutants were obtained with *OsBZR2*, suggesting that those genotypes resulted in lethality and this gene encodes the dominant BES1/BZR1 transcription factor in rice. It is perhaps surprising that *Osbzr2* was the only single mutant to have a phenotype, as the two maize single mutants that exhibited mutant phenotypes (Wang et al., 2022) are more closely related to the three other rice genes (Figure 11).

The rice U-box E3 ligase, *Os*PUB24, was shown to ubiquitinate *Os*BZR1 and result in the subsequent degradation of *Os*BZR1 by the 26S proteasome (Min et al., 2019). Knock-out mutants of *Ospub24* were hypersensitive to BL treatment, had less upright leaves, and had increased seedling growth of both the root and shoot (Min et al., 2019). These results were similar to what was observed in Arabidopsis for *Atpub39*;*pub40*;*pub41* triple mutants which accumulated more *At*BZR1 in roots than their wild-type siblings. Consistent with this observation, overexpression of *Os*PUB40 resulted in lower *At*BZR1 levels in a root specific manner (Kim et al., 2019). No mutants of BES1/BZR1 have been described in Brachypodium or Setaria and given the conserved redundancy of gene function described in Arabidopsis, soybean, and rice, higher order mutant combinations will most likely be necessary to identify their function.

## Acknowledgements

Mention of trade names or commercial products in this publication was solely for the purpose of providing specific information and does not imply recommendation or endorsement by the U.S. Department of Agriculture. The U.S. Department of Agriculture is an equal opportunity provider and employer. This research used resources provided by the SCINet project of the USDA Agricultural Research Service, ARS project number 0500-00093-001-00-D. This work was supported by USDA Hatch project number IOW03649 (EV). This work was supported by the Department of Agriculture, National Institute of Food and Agriculture (USDA-NIFA) fellowship number #2019-67012-29655 (NBB).

## Author Contributions

BZ, EV, and NBB wrote the manuscript.

