## Supplemental information for "Conservation and diversification of genes regulating brassinosteroid biosynthesis and signaling"

### Material and Methods

#### *Phylogenetic analysis*

Gene IDs for Arabidopsis (version TAIR10) BR related genes were obtained from ThaleMine (version 5.0.2-20210109; (Krishnakumar et al., 2017). Sequence similarity between known Arabidopsis BR genes and maize, rice, Brachypodium, Setaria, soybean, and Physcomitrella genes was determined using Phytozome (version 12.1; (Goodstein et al., 2012). Protein sequences for Arabidopsis, soybean (version 2.1), rice (version IRGSP-1.0), Setaria (version 2.0), Brachypodium (version 3.0), and Physcomitrella (version 3.3) were downloaded from Phytozome. The BKI1 like genes in Physcomitrella did not have high sequence similarity, thus a homologous gene to BKI1 was identified in *Zostera marina* (version 2.2) and was used for the phylogenetic analysis. Maize protein sequences (version Zm-B73-REFERENCE-NAM-5.0) were downloaded from the Maize Genetics and Genomics Database (https://maizegdb.org; (Portwood et al., 2019).

Protein sequences for each gene family were aligned using ClustalW2 (Larkin et al., 2007) within the Mesquite software package (version 3.61; (Maddison and Maddison, 2019). Aligned sequences were outputted as fasta files and approximate-maximum-likelihood phylogenetic trees were created using FastTreeMP (version 2.1.8; (Price et al., 2010)) which was run on the USDA SciNet cluster. Maximum clade credibility (.tre) files were downloaded and visualized in Dendroscope (version 3.7.2; (Huson and Scornavacca, 2012)). Physcomitrella gene(s) were used as outgroups, except for the BKI1 family when *Zostera marina* was used, and all phylogenetic trees were rooted on these genes. Bootstrap values were represented at selected nodes to show branch strength.

#### *RNA transcript analysis and heatmap creation*

Raw RNA sequence reads were acquired as SRA files from the short-read archive (https://www.ncbi.nlm.nih.gov/sra) for maize, rice, and Setaria data. The Arabidopsis raw RNA sequence reads were downloaded as fastq files from the European nucleotide archive (https://www.ebi.ac.uk/ena/browser/view/). A description of the datasets with accession numbers used in this work is described in Supplemental Tables 1-6 (Severin et al., 2010; Davidson et al., 2012; Stelpflug et al., 2016; Zhu et al., 2018; Harrop et al., 2019; Mergner et al., 2020; Parvathaneni et al., 2020; Cheng et al., 2021). The SRA files for maize and rice data were converted to fastq files using sra-toolkit (version 2.10.9; http://ncbi.github.io/sra-tools/). The fastq files for all datasets were aligned to their most current respective genome using tophat2 (version 2.1.0; (Kim et al., 2013)) in conjunction with bowtie2 (version 2.3.4; (Langmead and Salzberg, 2012)) using default settings. The genome fasta files and respective gff3 files for maize (Zm-B73-REFERENCE-NAM-5.0), rice (IRGSP-1.0), Brachypodium (Brachypodium_distachyon_v3.0.56), Setaria (Setaria_viridis_v2.0), soybean (Glycine_max_v2.1.56) and Arabidopsis (TAIR10) were downloaded from EnsemblPlants (https://plants.ensembl.org; (Bolser et al., 2016)).

Aligned reads in the BAM file format were sorted using samtools (version 1.10; (Li et al., 2009)). Sorted reads were analyzed for Fragments Per Kilobase of transcript per Million mapped reads (FPKM) and Transcripts Per Million (TPM) values at each locus using stringtie2 (version 2.1.0; (Kovaka et al., 2019)). Outputs were obtained in the table format and merged into a single file for each species (Supplemental Files 1-4). For tissues with multiple sequence datasets from an individual species, the average FPKM and TPM values were calculated across the datasets, and this was used for the final expression values to create the heatmaps.

Heatmaps were created as a percentage for TPM values across tissues for each gene. The highest TPM value for each gene was given a value of 100 (dodgerblue) and the lowest was given a value of zero (yellow). To calculate the average expression for each gene to enable comparison of expression values between genes and species, the average FPKM value was calculated across all tissues analyzed for a respective gene in each species. The heatmap for average expression comparison for genes across all species was created as previously described above.

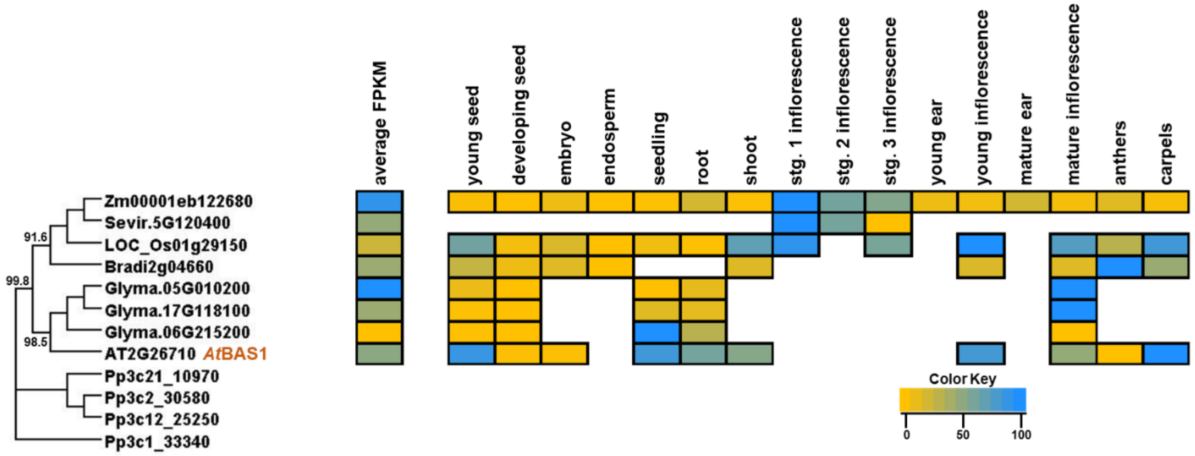


**Supplemental Figure 1. Phylogeny and transcript abundance across different tissues of PHYB ACTIVATION-TAGGED SUPPRESSOR1 (BAS1) family involved in brassinosteroid perception.**
Maximum-approximate-likelihood phylogenetic tree of BAS1 amino acid sequences from maize (Zm), Setaria (Sevir), rice (Os), Arabidopsis (AT), soybean (Glyma), Brachypodium (Bradi), and Physcomitrella (Pp) represented as the outgroup. The relative TPM values from re-analyzed publicly available datasets from different developmental tissues are represented as a heatmap depicted next to each respective gene within the family. The yellow color indicates the lowest individual transcript abundance in each tissue and the dodgerblue color indicates the highest abundance in a given tissue. To compare transcript abundance across genes and species within the family, the average FPKM was determined across all tissues analyzed and is presented in the average FPKM column (left) with the same color distribution as previously described. See Supplemental Tables 1-4 for description of tissue types.

***Supplemental Table 1.*** Description and sources of publicly available maize sequence data

| Tissue Type | Description | SRA accession files |
|---|---|---|
| young seed | whole seed 6 DAP[a] | SRR531904,SRR531905,SRR531906 |
| developing seed | whole seed 12 DAP[a] | SRR940283,SRR940284,SRR940286 |
| embryo | embryo 24 DAP[a] | SRR533835,SRR533845,SRR533848 |
| endosperm | endosperm 24 DAP[a] | SRR531946,SRR533828,SRR533829 |
| seedling | V3 topmost leaf[a] | SRR531218,SRR531219,SRR531220 |
| root | whole root system 7d[a] | SRR1620957,SRR1620953,SRR1620971 |
| shoot | V1 4d PE pooled leaves[a] | SRR531205,SRR531206,SRR531207, SRR531208,SRR940236,SRR531209, SRR531210 |
| stg. 1 inflorescence | 1-2 mm tassels[b] | SRR999038,SRR999039 |
| stg. 2 inflorescence | 2-4 mm tassels[b] | SRR999040,SRR999041 |
| stg. 3 inflorescence | 4-7 mm tassels[b] | SRR999042,SRR999043 |
| young ear | V18 immature cob[a] | SRR531886,SRR531887,SRR531888 |
| young inflorescence | V13 immature tassel[a] | SRR531878,SRR531879,SRR531880 |
| mature ear | pre-pollination cob R1[a] | SRR531889,SRR531890,SRR531891 |
| mature inflorescence | V18 meiotic tassel[a] | SRR940258,SRR531882,SRR531883, SRR531884 |
| anthers | anthers R1[a] | SRR531895,SRR531896,SRR531897 |
| carpels | silks R1[a] | SRR531892,SRR531893,SRR531894 |

[a]Stelpflug et al. 2015
[b]Parvathaneni et al. 2020

***Supplemental Table 2.*** Description and sources of publicly available Setaria sequence data

| Tissue Type | Description | SRA accession files |
|---|---|---|
| young seed | n/a | |
| developing seed | n/a | |
| embryo | n/a | |
| endosperm | n/a | |
| seedling | n/a | |
| root | n/a | |
| shoot | n/a | |
| stg. 1 inflorescence | 12 DAS A10 inflorescence[a] | SRR7704031,SRR7704032,SRR7704033 |
| stg. 2 inflorescence | 14 DAS A10 inflorescence[a] | SRR7704034,SRR7704035,SRR7704036, SRR7704037 |
| stg. 3 inflorescence | 16 DAS A10 inflorescence[a] | SRR7704042,SRR7704043,SRR7704044, SRR7704045 |
| young ear | n/a | |
| young inflorescence | n/a | |
| mature ear | n/a | |
| mature inflorescence | n/a | |
| anthers | n/a | |
| carpels | n/a | |

[a]Zhu et al. 2018

***Supplemental Table 3.*** Description and sources of publicly available rice sequence data

| Tissue Type | Description | SRA accession files |
|---|---|---|
| young seed | seed 5 d after pollination[a] | SRR352194 |
| developing seed | seed 10 d after pollination[a] | SRR352207 |
| embryo | embryo 25 d after pollination[a] | SRR352204 |
| endosperm | endosperm 25 d after pollination[a] | SRR352206,SRR352209 |
| seedling | seedling leaves 20 d after sowing[a] | SRR352184,SRR352211 |
| root | 21 d old seedling roots[b] | SRR12011286,SRR12011285, SRR12011284 |
| shoot | seedling shoots[a] | SRR042529 |
| stg. 1 inflorescence | indeterminate stage[c] | SRR8517040,SRR8517041,SRR8517042 |
| stg. 2 inflorescence | n/a | |
| stg. 3 inflorescence | determinate stage[c] | SRR8517039,SRR8517043,SRR8517044 |
| young ear | n/a | |
| young inflorescence | inflorescence 10 d before emergence[a] | SRR352189 |
| mature ear | n/a | |
| mature inflorescence | inflorescence after emergence[a] | SRR352187 |
| anthers | whole anther at anthesis[a] | SRR352190 |
| carpels | whole pistil at anthesis[a] | SRR352192 |

[a]Davidson et al. 2012
[b]Cheng et al. 2021
[c]Harrop et al. 2019

***Supplemental Table 4.*** Description and sources of publicly available Arabidopsis sequence data

| Tissue Type | Description | Run accession files |
|---|---|---|
| young seed | silique stage 4[a] | ERR3333434 |
| developing seed | embryo stage 6[a] | ERR3333436 |
| embryo | embryo stage 8[a] | ERR3333438 |
| endosperm | n/a | |
| seedling | rosette leaf 12 -22 d[a] | ERR3333425 |
| root | 7 d root tip and upper zone averaged[a] | ERR3333408,ERR3333409 |
| shoot | 7 d mixed shoot apical meristem, cotyledon, and first leaves[a] | ERR3333411 |
| stg. 1 inflorescence | n/a | |
| stg. 2 inflorescence | n/a | |
| stg. 3 inflorescence | n/a | |
| young ear | n/a | |
| young inflorescence | flower stage 9[a] | ERR3333426 |
| mature ear | n/a | |
| mature inflorescence | flower stage 13[a] | ERR3333430 |
| anthers | stamen- flower stage 15[a] | ERR3333410 |
| carpels | carpel- flower stage 15[a] | ERR3333421 |

[a]Mergner et al. 2020

***Supplemental Table 5.*** Description and sources of publicly available soybean sequence data

| Tissue Type | Description | Run accession files |
|---|---|---|
| young seed | seed 14-17 d after fertilization[a] | Soy:001290 |
| developing seed | seed 21 d after fertilization[a] | Soy:001291 |
| embryo | n/a | |
| endosperm | n/a | |
| seedling | young leaf[a] | Soy:000252 |
| root | root structures[a] | Soy:001183 |
| shoot | n/a | |
| stg. 1 inflorescence | n/a | |
| stg. 2 inflorescence | n/a | |
| stg. 3 inflorescence | n/a | |
| young ear | n/a | |
| young inflorescence | n/a | |
| mature ear | n/a | |
| mature inflorescence | open flower[a] | Soy:001277 |
| anthers | n/a | |
| carpels | n/a | |

[a]Severin et al. 2010

***Supplemental Table 6.*** Description and sources of publicly available Brachypodium sequence data

| Tissue Type | Description | Run accession files |
|---|---|---|
| young seed | seed 5 d after pollination[a] | SRR352139 |
| developing seed | seed 10 d after pollination[a] | SRR352141 |
| embryo | embryo 25 d after pollination[a] | SRR352138,SRR352144 |
| endosperm | endosperom 25 d after pollination[a] | SRR352142, SRR352143 |
| seedling | leaf 20 d after sowing[a] | SRR349785,SRR352143 |
| root | n/a | |
| shoot | n/a | |
| stg. 1 inflorescence | n/a | |
| stg. 2 inflorescence | n/a | |
| stg. 3 inflorescence | n/a | |
| young ear | n/a | |
| young inflorescence | inflorescence 10 d before emergence[a] | SRR349786 |
| mature ear | n/a | |
| mature inflorescence | inflorescence at emergence[a] | SRR349787 |
| anthers | anther at anthesis[a] | SRR352140 |
| carpels | pistil at anthesis[a] | SRR352137 |

[a]Davidson et al. 2012
